# Cysteine crosslinking in native membranes establishes the transmembrane architecture of Ire1

**DOI:** 10.1101/772772

**Authors:** Kristina Vaeth, Carsten Mattes, John Reinhard, Roberto Covino, Heike Stumpf, Gerhard Hummer, Robert Ernst

## Abstract

The endoplasmic reticulum (ER) is a key organelle of membrane biogenesis and crucial for the folding of both membrane and secretory proteins. Sensors of the unfolded protein response (UPR) monitor the unfolded protein load in the ER and convey effector functions for maintaining ER homeostasis. Aberrant compositions of the ER membrane, referred to as lipid bilayer stress, are equally potent activators of the UPR. How the distinct signals from lipid bilayer stress and unfolded proteins are processed by the conserved UPR transducer Ire1 remains unknown. Here, we have generated a functional, cysteine-less variant of Ire1 and performed systematic cysteine crosslinking experiments in native membranes to establish its transmembrane architecture in signaling-active clusters. We show that the transmembrane helices of two neighboring Ire1 molecules adopt an X-shaped configuration independent of the primary cause for ER stress. This suggests that different forms of stress converge in a common, signaling-active transmembrane architecture of Ire1.

**Summary:** The endoplasmic reticulum (ER) is a hotspot of lipid biosynthesis and crucial for the folding of membrane and secretory proteins. The unfolded protein response (UPR) controls the size and folding capacity of the ER. The conserved UPR transducer Ire1 senses both unfolded proteins and aberrant lipid compositions to mount adaptive responses. Using a biochemical assay to study Ire1 in signaling-active clusters, Väth *et al*. provide evidence that the neighboring transmembrane helices of clustered Ire1 form an ‘X’ irrespectively of the primary cause of ER stress. Hence, different forms of ER stress converge in a common, signaling-active transmembrane architecture of Ire1.

## Introduction

The endoplasmic reticulum (ER) marks the entry-point to the secretory pathway for soluble and membrane proteins. Under adverse conditions, accumulation of unfolded proteins causes ER stress and initiates the unfolded protein response (UPR). The UPR is mediated by the <u>I</u>nositol-<u>re</u>quiring enzyme 1 (Ire1) in budding yeast, and by the troika of IRE1α, the <u>P</u>KR-like <u>E</u>ndoplasmic <u>R</u>eticulum <u>K</u>inase (PERK), and the activating transcription factor 6 (ATF6) in vertebrates (Walter and Ron, 2011). Once activated, the UPR downregulates the production of most proteins and initiates a wide transcriptional program to upregulate ER chaperones, ER-associated degradation (ERAD), and lipid biosynthesis (Travers et al., 2000). Through these mechanisms, the UPR is centrally involved in cell fate decisions between life, death, and differentiation (Hetz, 2012). Insulin-producing β-cells, for example, rely on UPR signals for their differentiation into professional secretory cells, while chronic ER stress caused by an excess of saturated fatty acids kills them (Fonseca et al., 2009). Consistent with its broad effector functions, the UPR is associated with numerous diseases including diabetes, cancer, and neurodegeneration (Kaufman, 2002).

Ire1 is highly conserved among eukaryotes and represents the only transducer of ER stress in budding yeast (Nikawa and Yamashita, 1992; Kimata and Kohno, 2011). It is a type I transmembrane protein equipped with an ER-luminal sensor domain and two cytosolic effector domains: a kinase and an endoribonuclease (RNase) (Cox et al., 1993; Sidrauski and Walter, 1997; Mori et al., 1993). How exactly unfolded proteins activate the UPR via direct and indirect mechanisms is a matter of active debate (Karagöz et al., 2017; Gardner and Walter, 2011; Adams et al., 2019; Amin-Wetzel et al., 2017; Le and Kimata, 2021). ER stress caused by the accumulation of unfolded proteins leads to the oligomerization of Ire1 (Kimata et al., 2007), which activates the cytosolic effector kinase and RNase domains (Korennykh et al., 2009). The unconventional splicing of the *HAC1* precursor mRNA initiated by the RNase domain facilitates the production of an active transcription factor that controls a broad spectrum of genes with unfolded protein response elements (UPRE) in their promotor regions (Travers et al., 2000; Mori et al., 1992). A regulated *IRE1*-dependent decay of mRNA (RIDD) has been suggested as a parallel mechanism to reduce the folding load of the ER. However, RIDD does not seem to play the same important role in *Saccharomyces cerevisiae* as it does in *Saccharomyces pombe* or mammalian cells (Travers et al., 2000; Hollien and Weissman, 2006; Frost et al., 2012; Tam et al., 2014; Li et al., 2018).

Lipid bilayer stress due to aberrant compositions of the ER membrane is equally potent in activating the UPR (Promlek et al., 2011; Volmer et al., 2013; Surma et al., 2013). This membrane-based mechanism is conserved throughout evolution (Ho et al., 2018; Hou et al., 2014; Volmer et al., 2013) and has been associated with pathogenesis of type II diabetes and the lipotoxicity associated with obesity (Fonseca et al., 2009; Pineau and Ferreira, 2010). We have shown that Ire1 from baker’s yeast inserts an amphipathic helix (AH) into the luminal leaflet of the ER-membrane, thereby forcing the short, adjacent transmembrane helix (TMH) to tilt, which locally squeezes the bilayer (Halbleib et al., 2017). Aberrant stiffening of the ER membrane during lipid bilayer stress increases the free energy penalty for membrane deformations, thereby stabilizing oligomeric assemblies of Ire1 via a membrane-based mechanism (Halbleib et al., 2017; Ernst et al., 2018). Even though it is well-established that proteotoxic and lipid bilayer stress lead to the formation of Ire1 clusters (Kimata et al., 2007; Halbleib et al., 2017; Li et al., 2010; Belyy et al., 2020), it remains unexplored if these forms of ER stress have a distinct impact on the architecture of Ire1 within these clusters. It has been speculated that different forms of ER stress might induce conformational changes in the transmembrane region thereby allowing Ire1/IRE1α to mount custom-tailored adaptive programs (Hetz et al., 2020; Cho et al., 2019; Ho et al., 2020).

Here, we report on a systematic dissection of Ire1’s TMH region in signaling-active clusters. We have engineered a cysteine-less variant for a genomic integration at the endogenous *IRE1* locus and generated a series of constructs featuring single cysteines in the TMH region. This enabled us to develop a crosslinking approach and to study the transmembrane configuration of Ire1 in the natural environment of ER-derived membrane vesicles featuring a native complexity of lipids and proteins. This approach uncovers the overall transmembrane architecture of Ire1 and suggests an X-shaped configuration of the TMHs of neighboring Ire1 molecules. Our findings underscore the crucial importance of Ire1’s highly bent configuration in the TMH region for stabilizing an oligomeric state via a membrane-mediated mechanism. Most importantly, we provide direct evidence that proteotoxic and lipid bilayer stress converge in common architecture of the TMH region in signaling-active Ire1.

## Results

We used systematic cysteine-crosslinking in the TMH region of Ire1 to gain insight into the structural organization of signaling-active clusters during ER-stress. Recognizing that Ire1 is activated by aberrant physicochemical membrane properties (Halbleib et al., 2017; Ernst et al., 2018), which are exceedingly hard to mimic *in vitro*, we performed these experiments with microsomes exhibiting the natural complexity of ER proteins and lipids.

### Cysteine-less Ire1 is functional

We have generated a cysteine-less version of Ire1 that allows us to introduce single cysteine residues in the TMH region for subsequent crosslinking using copper sulfate (CuSO_4_). The cysteine-less construct is based on a previously established knock-in construct of *IRE1* that provides homogeneous, near-endogenous expression (Halbleib et al., 2017) and encodes for a fully-functional variant of Ire1 equipped with an 3xHA tag and a monomeric, yeast-enhanced GFP (yeGFP) inserted in a flexible loop at the position H875 (Fig. 1A) (van Anken et al., 2014; Halbleib et al., 2017). To generate a cysteine-less version, we substituted each of the twelve cysteines in the luminal, transmembrane and cytosolic domains with serine. Two cysteines in the signal sequence, which are co-translationally removed, remained in the final construct to ensure correct ER-targeting and membrane insertion (Fig. 1A). Cysteine 48 of yeGFP (C48^yeGFP^) was mutated to serine, while C70^yeGFP^ is present in the cysteine-less construct to ensure correct folding of the fluorescent protein (Costantini et al., 2015). Notably, C70^yeGFP^ is buried inside the green fluorescent protein (Ormö et al., 1996) and thus inaccessible for crosslinking agents under non-denaturing conditions.

**Figure 1.**
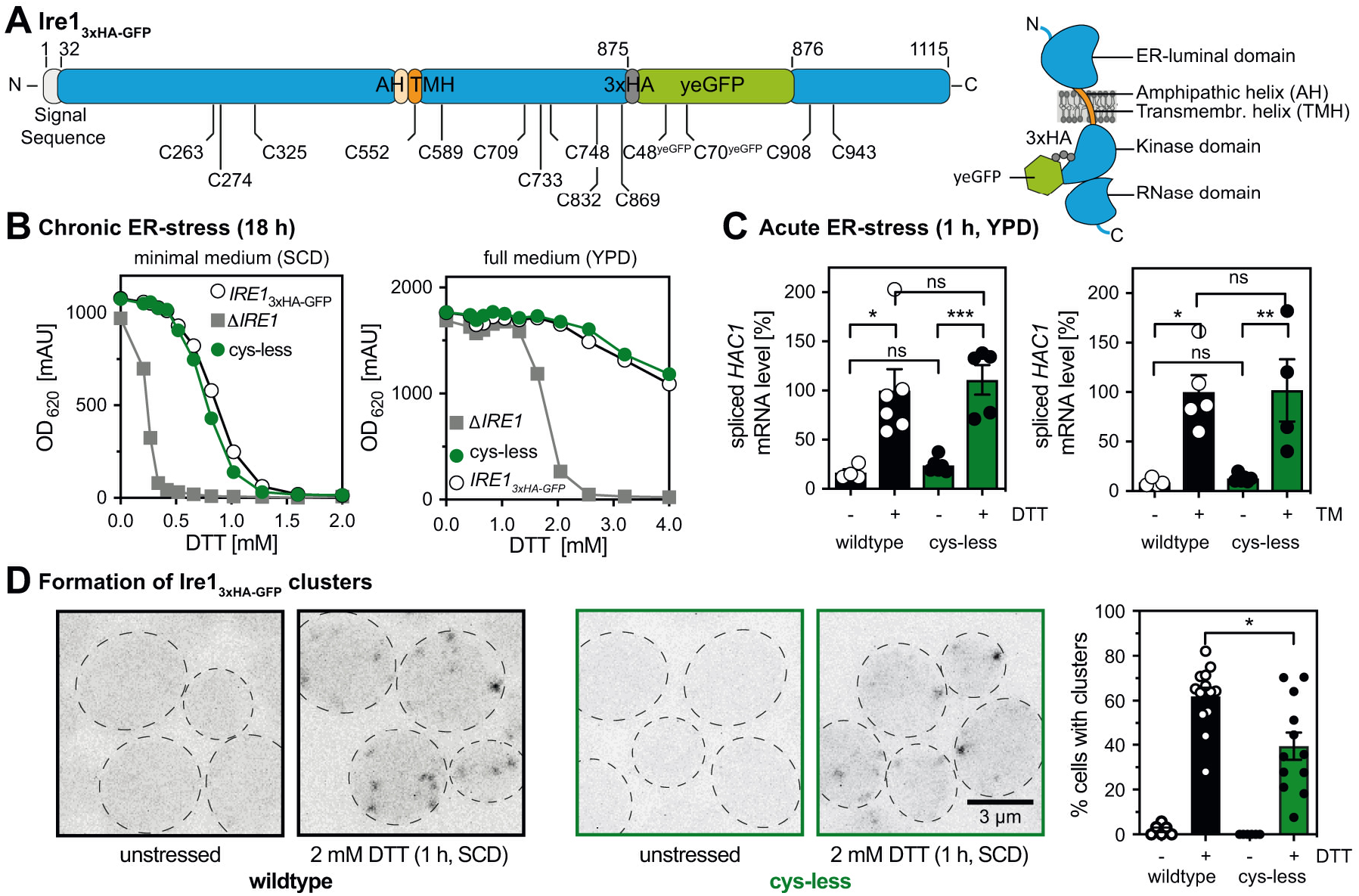
Cysteine-less Ire1 expressed from its endogenous locus is functional. (*A*) Schematic representations of the *IRE1*_3xHA-GFP_ construct indicating the position of cysteine residues and topology. All twelve cysteines of Ire1 and C48^yeGFP^ of yeGFP were substituted to serine to generate a cysteine-less variant. C70^yeGFP^ remains in the final construct. Two cysteines in the signal sequence of Ire1 are removed upon ER-translocation. (*B*) Resistance of the indicated strains to prolonged ER-stress. Stationary overnight cultures of the indicated strains were used to inoculate a fresh culture in full or minimal media to an OD_600_ of 0.2. After cultivation for 5 to 7 h at 30°C the cells were diluted with pre-warmed full or minimal media to an OD_600_ of 0.01. Cells were cultivated for 18 h at 30°C in the indicated media and stressed with DTT. The density of the resulting culture was determined using the OD_620_ or OD_600_. (*C*) The relative level of the spliced *HAC1* mRNA was determined by RT-qPCR in unstressed and acutely stressed cells. Exponentially growing cells of the indicated strains were used to inoculate a fresh culture in YPD medium to an OD_600_ of 0.2. After cultivation to an OD_600_ of 0.7, the cells were stressed for 1 h with either 4 mM DTT (left panel) or 1.0 μg/ml Tunicamycin (TM, right panel). The data were normalized to the level of the spliced *HAC1* mRNA in DTT-stressed cells with the *IRE1*_3xHA-GFP_ wildtype construct. (*D*) Cells were cultivated from OD_600_ of 0.2 to OD_600_ of 0.7 in SCD medium and then either left untreated or stressed with 2 mM DTT for 1 h. Life cells were mounted on agar slides and z-stacks were recorded using confocal microscopy. Cells and clusters of Ire1 were automatically detected and quantified. All data are represented as the mean ± SEM of at least three independent experiments. Significance was tested by an unpaired, two-tailed Student’s t test, with the exception of (C), which was analyzed using a Kolmogorov–Smirnov test. ***p<0.001, **p<0.01, *p<0.05, ns: not significant.

The steady-state levels of wildtype and cysteine-less Ire1 are comparable (*Suppl. Materials* Fig. S1A). Cysteine-less Ire1 is properly integrated into the membrane as shown by subcellular fractionation (*Suppl. Materials* Fig. S1B) and extraction assays (*Suppl. Materials* Fig. S1C), thereby matching previous observations for wildtype Ire1 (Kimata et al., 2007; Halbleib et al., 2017). The functionality of cysteine-less Ire1 was analyzed using a sensitive assay scoring for the growth of cells exposed to inducers of ER stress (Halbleib et al., 2017). Liquid cultures in either minimal (synthetic complete dextrose; SCD) or full (yeast peptone dextrose; YPD) medium were exposed to different concentrations of the reducing agent DTT interfering with disulfide bridge formation in the ER. After 18 h of cultivation, the optical densities (OD) of these cultures were determined. Cells producing either wildtype or cysteine-less Ire1 are phenotypically indistinguishable by this assay and substantially more resistant to DTT than cells lacking *IRE1* (Fig. 1B). This suggests that cysteine-less Ire1 is functional and capable to mount an adaptive UPR.

The functionality of cysteine-less Ire1 was further validated by quantifying the mRNA levels of spliced *HAC1* (Fig. 1C) and the mRNA level of the UPR-target gene *PDI1* (*Suppl. Materials* Fig. S1D) in both stressed and unstressed cells. We used either DTT or Tunicamycin, an inhibitor of N-linked glycosylation, to induce proteotoxic stress for hour and analyzed lysates from stressed and unstressed cells by RT-qPCR. As expected, the level of the spliced *HAC1* mRNA was several-fold higher in stressed versus unstressed cells and this upregulation is observed in both wildtype and cysteine-less Ire1-producing cells. (Fig. 1C). Control experiments validated also a comparable degree of *HAC1* mRNA splicing in WT or cysteine-less Ire1 producing cells stressed with DTT (*Suppl. Materials* Fig. S1D). We also observed an upregulation of the *PDI1* mRNA in response to ER-stress, albeit to slightly lower extent for the cysteine-less version compared to the wildtype construct (*Suppl. Materials* Fig. S1E). Using confocal microscopy and by applying an automated pipeline to identify cells with and without fluorescent clusters, we show that both cysteine-less and wildtype Ire1 cluster under conditions of ER stress, but not in unstressed cells (Fig. 1D). Notably, confocal microscopy can only identify large clusters of Ire1, while dimers and smaller assemblies escape our detection. Furthermore, the detection of Ire1 in unstressed cells is particularly challenging in our case, because our knock-in strategy aims to provide a close-to-endogenous level of *IRE1* expression (Halbleib et al., 2017). This is important because even the mild degree of overexpression when using an endogenous promotor from a CEN-based plasmid (Karim et al., 2013) is likely to interfere with normal UPR function by favoring dimerization and oligomerization. Using our setup, we robustly detect GFP-positive clusters of Ire1 (Fig. 1D) in stressed cells, while the tendency of clustering is somewhat lower for the cysteine-less Ire1 compared to the wildtype (Fig. 1D). Colocalization of GFP-positive clusters with an ER-targeted variant of dsRed-HDEL confirms the ER localization of wildtype and cysteine-less Ire1 in DTT-stressed cells (*Suppl. Materials* Fig. S1F). In line with the functional data (Fig. 1B,C), we conclude that both wildtype and cysteine-less Ire1 can mount robust responses to acute and prolonged forms of ER stress.

### Crosslinking of Ire1’s TMH in ER-derived microsomes

We established a strategy to crosslink single-cysteine variants of Ire1 via copper sulfate (CuSO_4_) in microsomes derived from the ER of stressed cells (Fig. 2A-C). Our approach has several advantages over previous attempts: Ire1 is studied i) as a full-length protein, ii) at the near-endogenous level, iii) in its natural, complex membrane environment, iv) with a spatial resolution of one residues and v) in a signaling-active state. In contrast to mercury chloride (HgCl_2_), which crosslinks by forming covalent bonds with two nearby cysteines (Soskine et al., 2002), CuSO_4_ is ‘traceless’ by catalyzing the oxygen-dependent formation of a disulfide bond (Bass et al., 2007). We performed the X-linking experiments on ice and with CuSO_4_ (instead of the more reactive Cu^2+^-phenanthroline) to prevent the loss-of-signal from unspecific crosslinking and/or aggregation. Even though every crosslinking approach on membrane proteins faces that the challenge of varying efficiencies at different depths in the membrane, Cu^2+^-mediated crosslinking has been successfully used to interrogate and establish structure-function relationships of membrane proteins (Falke and Koshland, 1987; Bass et al., 2007; Matthews et al., 2011; Lopez-Redondo et al., 2018). Here, we have studied the configuration of Ire1’s TMH in UPR-signaling clusters, which are long-lived and stable for minutes (Kimata et al., 2007; Cohen et al., 2017). Because CuSO_4_-mediated crosslinking occurs on the same timescale, it can provide useful structural information even though it leads to the formation of covalent disulfide bonds under our experimental conditions.

**Figure 2.**
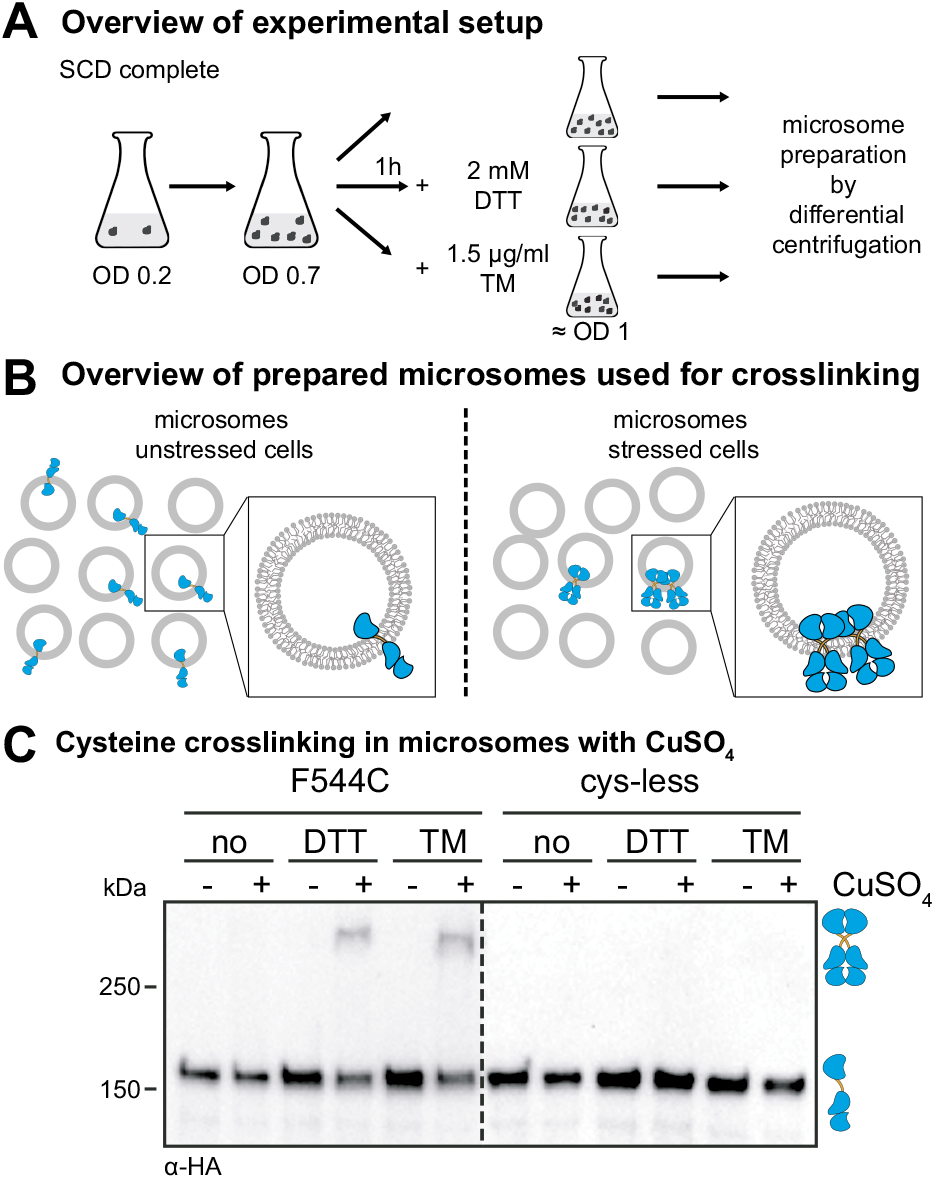
The crosslinking of Ire1 via single cysteines in microsomes requires CuSO4 and pre-formed clusters. (*A*) Cultivation of yeast cells for cysteine crosslinking. A culture in SCD medium was inoculated with stationary cells to an OD_600_ of 0.2. After cultivation at 30°C to an OD_600_ of 0.7, the clustering of Ire1 was induced either by DTT (1 h, 2 mM, SCD) or TM (1 h, 1.5 μg/ml, SCD) as indicated. After harvesting, the cells were lysed and used to prepare microsomes. (*B*) Schematic representation of the cysteine crosslinking with CuSO_4_. Only microsomes from stressed cells contain clusters of Ire1 clusters that can crosslinked via cysteines using CuSO_4_. (*C*) Crosslinking of a single-cysteine-variant of Ire1 in microsomes. The indicated strains were cultivated in the presence and absence of ER-stressors as described in (*A*). 80 OD equivalents of cells were harvested and microsomes were prepared. 8 μl microsomes (1 mg/ml protein) were mixed with 2μl of 50 mM CuSO_4_ and the sample was incubated on ice for 5 min to catalyze cysteine crosslinking. The reaction was stopped by the addition of 2 μl 1 M NEM, 2 μl 0.5 M EDTA and 4 μl membrane sample buffer. The resulting samples were analyzed by SDS-PAGE and immunoblotting using anti-HA antibodies.

Cells expressing either a cysteine-less variant of Ire1 or a variant with a single-cysteine in the TMH region (F544C) were cultivated to the mid-exponential phase in minimal medium (Fig. 2A). These cells were either left untreated or stressed for 1 h with either DTT (2 mM) or TM (1.5 μg/ml) to cause ER-stress, which leads to the formation of Ire1-clusters (Kimata et al., 2007; Halbleib et al., 2017; Belyy et al., 2020). We used such an early time-point to minimize the contribution of secondary effects from stress- and UPR-dependent reprogramming of the cell. We then isolated crude microsomes from these cells and incubated them on ice for 5 min either in the presence or absence of 10 mM CuSO_4_ to catalyze the formation of disulfide bonds by oxidizing nearby sulfhydryl groups (Kobashi, 1968). Given the low copy number of ~260 for Ire1 (Ghaemmaghami et al., 2003) and the fragmentation of the ER during microsome preparation, we expect to detect crosslinking of single-cysteine variants of Ire1 only when it was clustered prior to the preparation (Fig. 2B).

Immunoblotting of the resulting samples revealed a prominent, HA-positive signal corresponding to monomeric Ire1 and a less-pronounced HA-positive signal from a band with lower electrophoretic mobility that was only observed when i) Ire1 contained a single-cysteine in the TMH region (F544C), ii) the microsomes were prepared from stressed cells (either DTT or TM), and iii) when crosslinking was facilitated by CuSO_4_ (Fig. 2C). This suggests a remarkably specific formation of covalent, disulfide bonds between two Ire1 molecules in the TMH region, despite the presence of numerous other, potentially competing membrane proteins with exposed cysteines in the ER. The observed degree of crosslinking was somewhat low considering that up to 70-85% of Ire1 may reside in signaling-active clusters under conditions of ER stress (Aragón et al., 2009). For our crosslinking approach, however, we used a slightly milder condition to induce ER stress (2 mM DTT instead of 10 mM) and performed all experiments with an *IRE1* knock-in strain that provides a more native-like expression level (Halbleib et al., 2017; Aragón et al., 2009). Notably, the signal from the crosslinked species was neither increased by the use of more reactive crosslinking agents (e.g. HgCl_2_ or Cu^2+^-phenanthroline) nor by harsher crosslinking conditions (higher temperatures or increased concentrations of the crosslinking agent). In fact, more reactive agents and harsher conditions only caused a loss of the total HA-positive signal presumably due to an unspecific crosslinking and/or aggregation of Ire1 (*data not shown*). A Co-IP analysis using Flag- and HA-tagged Ire1 variants produced in the same cell and crosslinked in microsomes via the native cysteine (C552) verified that the additional band with low electrophoretic mobility represents disulfide-linked, SDS-resistant dimers of Ire1 (*Suppl. Materials* Fig. S2A). In fact, treating a crosslinked species of Ire1 with heat under reducing conditions revealed full reversibility of disulfide bond formation (*Suppl. Materials* Fig. S2B). We conclude that CuSO_4_ can catalyze the formation of disulfide bridges between two neighboring Ire1 molecules, when they are present in pre-formed clusters and isolated in microsomes from stressed cells.

### A crosslinking screen in the TMH region of Ire1

Next, we generated a set of thirteen mutant variants of Ire1 each containing a single cysteine in the TMH region starting with E540C at the transition between the AH and the TMH (Fig. 3A) and ending at the native C552, which is substituted to serine in cysteine-less Ire1. Our scanning approach covered more than three helical turns and almost the entire short TMH of Ire1 (Fig. 3A,B). Systematic crosslinking of these variants can provide important insight into the organization of Ire1’s TMH in signaling-active clusters. An important prerequisite for a structural interpretation is that the single-cysteine substitutions required to form the crosslinks do neither affect the oligomerization nor the activity of Ire1.

**Figure 3.**
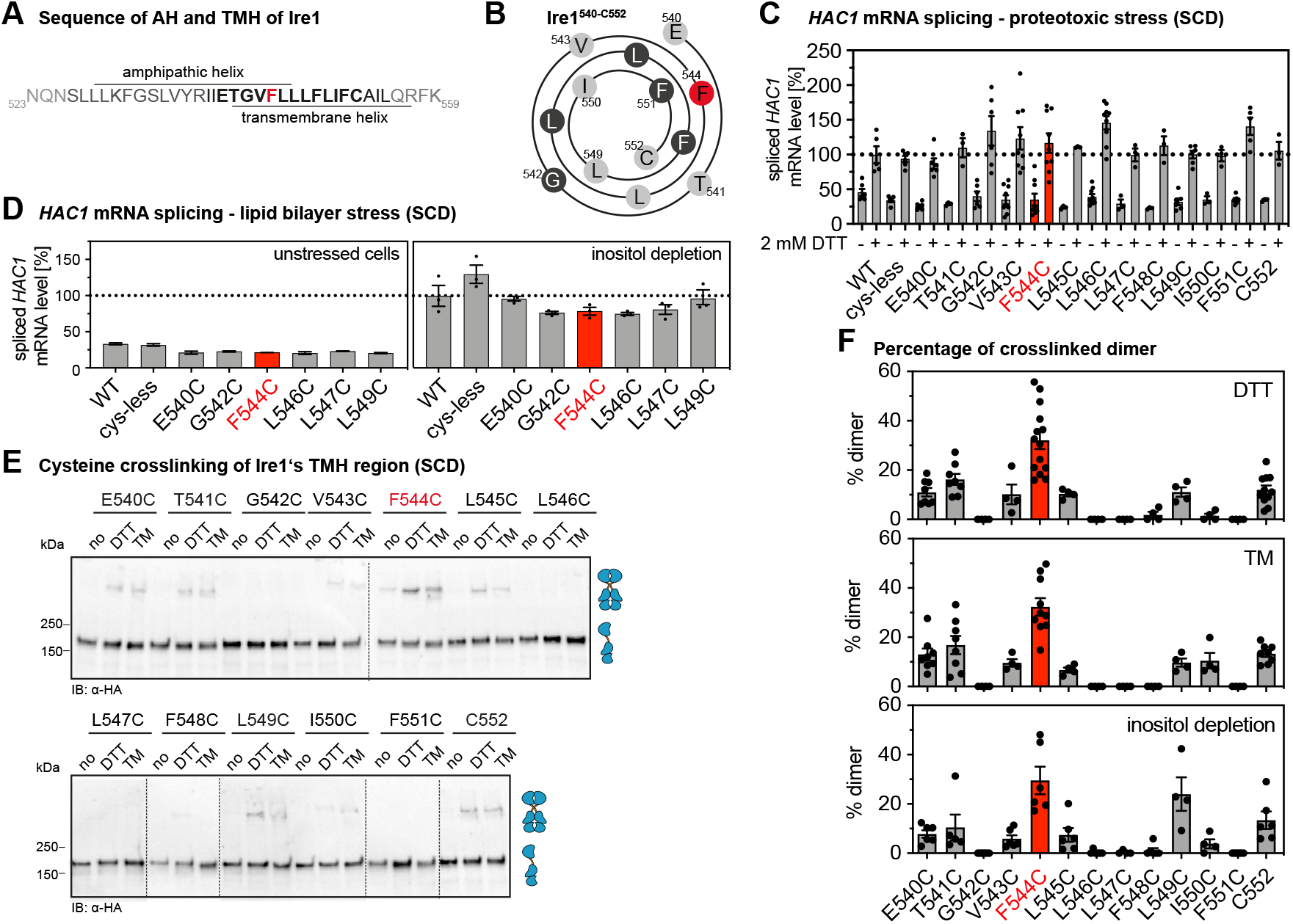
Systematic crosslinking of cysteines in the TMH region of Ire1 reveals a specific configuration during ER stress. (*A*) Primary structure of ER-luminal AH of Ire1 and the short TMH. Almost every residue of the short TMH (shown in bold) was substituted individually by cysteine for the cysteine crosslinking strategy. (*B*) Helical wheel representation of Ire1’s TMH (Ire1^540-552^). (*C*) The level of the spliced *HAC1* mRNA was determined from the indicated strains by RT-qPCR for either unstressed cells or cells stressed with 2 mM DTT for 1 h (for details see Fig. 3E). The data are normalized to the level of the spliced *HAC1* mRNA in stressed cells with a tagged, wildtype variant of Ire1. (*D*) The level of the spliced *HAC1* mRNA was determined from the indicated strains by qPCR using stressed (inositol-depleted) and unstressed cells. The data are normalized to the level of the spliced *HAC1* mRNA splicing caused by 2 mM DTT, as determined in (C). (*E*) A culture in SCD medium was inoculated with stationary cells to an OD_600_ of 0.2. After cultivation at 30°C to an OD_600_ of 0.7, Ire1-clustering was induced either by DTT (1 h, 2 mM, SCD) or TM (1 h, 1.5 μg/ml, SCD). 8 μl microsomes (1 mg/ml protein) from unstressed (no) and stressed cells were mixed with 2μl of 50 mM CuSO_4_ and the sample was incubated on ice for 5 min to catalyze cysteine crosslinking. The reaction was stopped, and the sample was analyzed by SDS-PAGE and immunoblotting using anti-HA antibodies. (*F*) Quantification of cysteine-crosslinking of the indicated variants of Ire1 in microsomes isolated cells stressed either by DTT, TM, or inositol-depletion. Cells were cultivated and treated as described in (E). For inositol depletion, a culture was inoculated with exponentially growing cells to an OD_600_ of 0.5 and cultivated for 3 h at 30°C in inositol-free medium (a representative immunoblot after crosslinking is shown in *Suppl. Materials* Fig. S3B. The percentage of crosslinked species was determined by densitometry. Data are represented as the mean ± SEM of at least three independent experiments.

We therefore subjected all Ire1 variants with engineered cysteine residues (E540C to F551C) to a sensitive, cell-based assay to ascertain the functionality of the UPR under conditions of prolonged ER stress (*Suppl. Materials* Fig. S3A). Consistent with the functional role of the AH adjacent to the short TMH (Halbleib et al., 2017), we found that the substitution of AH-residues to cysteine (E540C, T541C, or G542C) impaired the response to ER stress as evident from an increased sensitivity of the respective cells to DTT (*Suppl. Materials* Fig. S3A). The substitution of TMH residues (V543C-F551C), by contrast, did not cause any apparent functional defect (*Suppl. Materials* Fig. S3A). Hence, these TMH variants are suitable to map the transmembrane architecture via cysteine crosslinking. In order to validate the functionality of these variants with a more direct assay, we systematically quantified the level of the spliced *HAC1* mRNA in stressed and unstressed cells both under conditions of proteotoxic (Fig. 3C) and lipid bilayer stress (Fig. 3D), which is caused by inositol-depletion (Promlek et al., 2011; Surma et al., 2013). Because these data are normalized to the level of the spliced *HAC1* mRNA in DTT-stressed cells, it is possible to compare the UPR activity between these conditions (Fig. 3C,D). We find a similar level of the *HAC1* mRNA in stressed cells and, consistently, a comparable degree of *HAC1* mRNA splicing in cells by either DTT or inositol-depletion (*Suppl. Materials* Fig. S3B). All single-cysteine variants were functional and responsive to proteotoxic stress (Fig. 3C). Likewise, the subset of variants tested under conditions of lipid bilayer stress showed robust activation of the UPR (Fig. 3D). Because the steady-state level of all Ire1 variants was also comparable (*Suppl. Materials* Fig. S3C), we could proceed with mapping the TMH region.

We subjected the entire set of single cysteine variants to the cysteine-crosslinking procedure (Fig. 3E; *Suppl. Materials* Fig. S3D) and determined the fraction of crosslinked Ire1 for construct (Fig. 3F). While some variants (e.g. G542C or L546C) showed no detectable crosslinking, a significant portion of them (e.g. T541C or L549C) could be crosslinked under the given experimental conditions (Fig. 3E, F). The F544C variant consistently exhibited the highest crosslinking efficiency (Fig. 3F). Notably, the differences in crosslinking are not caused by an aberrant oligomerization of Ire1, because confocal microscopy experiments with cells cultivated and treated as in the crosslinking experiments demonstrate the same degree of cluster formation of all single-cysteine variants upon ER stress as judged cluster size and intensity and compared to cysteine-less Ire1 (*Suppl. Materials* Fig. S3D-F).

### Different forms of ER-stress converge in a common architecture of the TMH region

Using the crosslinking assay, we could show that the overall pattern of crosslinking residues was independent of the condition of ER stress (Fig. 3F). Lipid bilayer stress and proteotoxic stress induced by either DTT or TM show essentially the same crosslinking pattern (Fig. 3F). These data strongly suggest that the overall structural organization of Ire1 is similar for different types of stress, at least in the TMH region. Notably, the L549C mutant showed significant crosslinking in cells stressed by DTT or TM, but even more during inositol-depletion (Fig. 3F). Because F544C, the best-crosslinking residue, and L549C seemingly lie on opposing sites of Ire1’s TMH as judged from a helical wheel representation (Fig. 3B), this raises the question if the corresponding residues in the native TMH can face each other at a low distance and at the same time. This point was addressed by molecular dynamics (MD) simulations further below.

Cysteine crosslinking can be used to infer structural models. The observed pattern of crosslinking residues in the TMH of Ire1 is very distinct to those observed in the TMH of the growth hormone receptor (Brooks et al., 2014) and the thrombopoietin receptor (Matthews et al., 2011), which form parallel dimers leading to a helical periodicity of crosslinking. Instead, our crosslinking data suggest an X-shaped configuration of the TMHs with the best-crosslinking residue F554 positioned at the crossing point. Intriguingly, such an arrangement would be consistent with the previously reported, highly tilted orientation of the monomeric TMH of Ire1, which is enforced by the adjacent, ER-luminal AH (Halbleib et al., 2017). However, it is important to realize that crosslinks might occur either within dimers of Ire1 or across dimers in higher oligomeric assemblies.

In order to obtain a structural representation, we used an experimentally validated model of the monomeric TMH region of Ire1 (Halbleib et al., 2017), generated a model of the dimer based on extensive molecular dynamics (MD) simulations in lipid membranes and integrated the crosslinking data with a particular attention on the contact between the two F544 (Fig. 4; *Suppl. Materials* Movie S1), which were restrained to face each other. The resulting model of the dimeric TMH region highlighted an highly bent configuration of each protomer leading to an X-shaped configuration of the dimer (Fig. 4; *Suppl. Materials* Movie S1). A substantial membrane thinning (Fig. 4B) and water penetration around the dimeric TMH region of Ire1 became apparent (*Suppl. Materials* Fig. S4A, Movie S1). It is tempting to speculate that this substantial degree of membrane deformation facilitates the access of Cu^2+^ ions to F544C for mediating efficient crosslinking (Fig. 3E, F). A thorough inspection of the trajectories revealed that the residues at position F544 and L549 can face their counterpart in an X-shaped dimer at the same time (*Suppl. Materials* Fig. S4C) thereby rendering the corresponding single-cysteine variants capable to crosslink (Fig. 3B). This would be unlikely if the TMHs would associate in a strictly parallel fashion. Inspecting the dynamics of Ire1’s TMH region in a MD simulation over a period of 1000 ns (*Suppl. Materials* Movie S1) underscored the stability of the overall X-shaped configuration, which nevertheless allowed for significant relative motions of the TMHs. In summary, our combined approach of biochemical crosslinking and MD simulations established a surprising configuration of Ire1’s TMH region with a particularly small interface between the TMHs.

**Figure 4.**
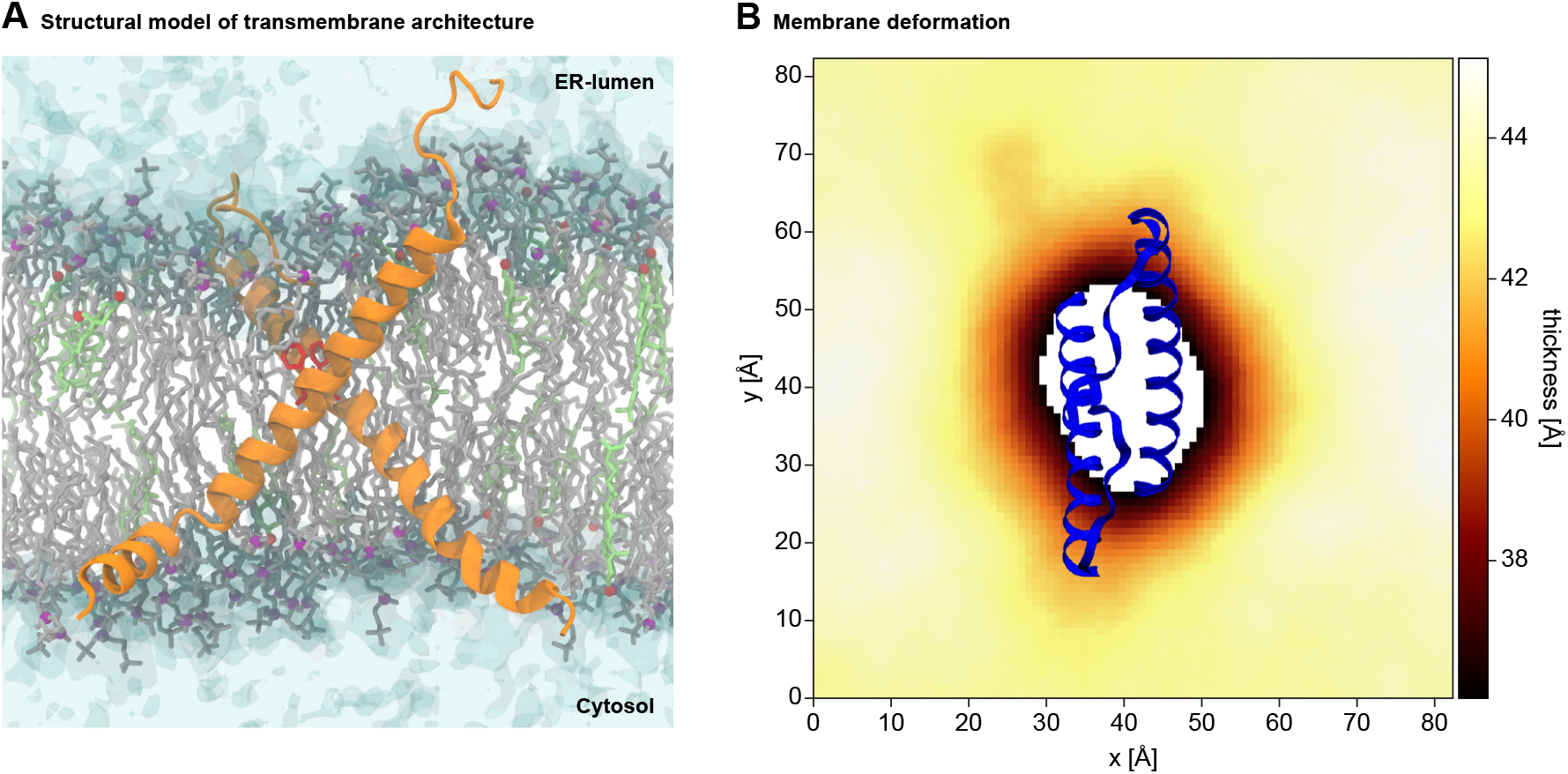
Structural model of the TMH region of Ire1. (*A*) Configuration of a model TMH dimer obtained from atomistic MD simulations. Protomers are shown as an orange ribbon with the two F544 residues highlighted in red. POPC lipids and their phosphate moieties are shown in gray and purple, respectively. Cholesterol molecules and their hydroxyl groups are shown in light green and red, respectively. Water is shown with a transparent surface representation. (*B*) Membrane thickness around the sensor peptide, defined as the average vertical distance between the two phosphate layers, averaged over MD simulations in POPE. A representative structure of the dimeric TMH region is shown in blue. For the standard error of the mean of the thickness profile see *Suppl. Materials* Fig. S4C.

### Validating the structural model of the TMH region of Ire1

Our crosslinking approach indicates that the TMH residue F544 is part of a small interface between Ire1 protomers, which might stabilize the unusual X-shaped transmembrane configuration of Ire1. Aromatic residues TMH residues have been implicated in sensing lipid saturation by Mga2 (W1042) (Covino et al., 2016; Ballweg et al., 2020) and lipid bilayer stress by the mammalian IRE1α (W547) (Cho et al., 2019). Despite a different position within the ER membrane, we wanted to test a similar role for F544 in Ire1 from baker’s yeast. We generated a F544A variant of Ire1, which contained the native C552 in the TMH as the only accessible residue for Cu^2+^-mediated crosslinking. A cell-based assay revealed that the F544A mutant was phenotypically indistinguishable from cysteine-less Ire1 (Fig. 5A) and the F544C mutant (Fig. 3C; *Suppl. Materials* Fig. S1A). This finding was corroborated by Cu^2+^-mediated crosslinking of C552 in microsomes isolated from stressed cells (DTT or TM). The intensity of the band corresponding to crosslinked Ire1 was unaffected by the F544A mutation (Fig. 5B). Thus, F544 does not contribute to the stability of Ire1 dimers and oligomers even though it is placed near to the equivalent residue on the opposing Ire1 protomer.

**Figure 5.**
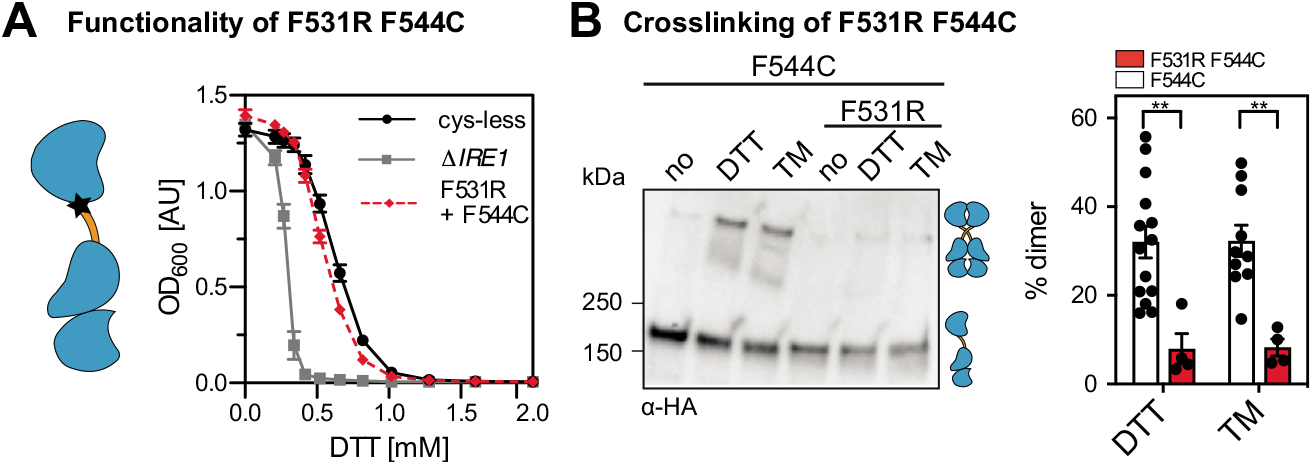
The impact of mutations in the TMH and the AH of Ire1 on its functionality and crosslinking propensity. (*A*) The ER-stress resistance of cells expressing the F544A variant of Ire1_3xHA-GFP_ containing the native cysteine 552 was determined. Stationary overnight cultures of the indicated strains were used to inoculate a fresh culture minimal media to an OD600 of 0.2. After cultivation for 5 to 7 h at 30°C, the cells were diluted in 96-well plates to an OD_600_ = 0.01 with pre-warmed minimal medium and cultivated in the presence of the indicated concentrations of DTT for 18 h at 30°C. The density of the resulting culture was determined using the OD_600_. (*B*) The impact of the F544A mutation on Ire1 degree of crosslinking via cysteine 552 was analyzed. The indicated strains were subjected to the crosslinking procedure as outlined in Fig. 3E. Data for the C552 variant are identical with the data in Fig. 3F. (*C*) ER-stress resistance of indicated cells including a single-cysteine variant (C552) of Ire1_3xHA-GFP_ with an AH-disrupting F531R mutation was determined. The cells were cultivated and treated as in (*A*). (*D*) The impact of the AH-disrupting F531R mutation on Ire1-crosslinking via cysteine 552 was determined. The indicated strains were subjected to the same crosslinking procedure as used for Fig. 3E. Data for the C552 single-cysteine variant are identical with the data in Fig. 3F. All data are represented as the mean ± SEM derived from at least three independent experiments. Significance was tested by an unpaired, two-tailed Student’s t test. *p<0.05, ns: not significant.

Previously, we have proposed that a tilted configuration of the monomeric TMH region, which is stabilized by a proximal AH, facilitates Ire1 to sense aberrant membrane properties (Halbleib et al., 2017; Covino et al., 2018). In fact, disrupting the amphipathic character of the AH by an F531R mutation increases the cellular sensitivity to ER stress (Fig. 5C) and reduces the crosslinking propensity via the native C552 residue in the TMH (Fig. 5D). These findings provide biochemical evidence that the AH contributes to the stability of either dimeric or oligomeric forms of Ire1, which are challenging to distinguish.

Similarly, when the AH-disrupting mutation F531R was combined with the F544C mutation (at the crossing-point of the X-shaped TMH-dimer), we observed only a very mild, yet significant functional defect (*Suppl. Materials* Fig. S5A) and a strongly reduced crosslinking propensity (*Suppl. Materials* Fig. S5B). This robust resistance to DTT is somewhat surprising considering the strongly reduced crosslinking propensity. However, the disruption of the AH changes the placement of the TMH in the membrane and the degree of membrane thinning and water penetration (Halbleib et al., 2017). We speculate that these combined changes would place the polar F544C residue more deeply in the hydrophobic core of the membrane, thereby affecting its propensity to undergo a Cu^2+^-catalyzed crosslinking, but at the same time favoring Ire1 dimerization. Notably, the F544C mutation alone does not lead to an increased UPR activity and ER stress resistance (Fig. 3C, D; *Suppl. Materials* Fig. S3A). In fact, the primary sequence of Ire1’s TMH can be systematically mutated (Fig. 3C; *Suppl. Materials* Fig. S3A), scrambled (in the case of the mammalian IRE1α) or exchanged altogether (Halbleib et al., 2017; Volmer et al., 2013) without causing a detectable functional defect. It therefore seems that a suitably placed polar residue in the TMH, here through the F544C mutation, becomes phenotypically relevant only when Ire1 is otherwise compromised. Beyond that, our data suggest that the overall architecture of the TMH region with an intact AH is relevant for normal UPR function.

### The TMH region of Ire1 makes dimer- and oligomer-specific contacts

Does the crosslinking of engineered cysteines in the TMH occur only within dimers of Ire1 or also across dimers in signaling-active clusters? The X-ray structure of the core ER-luminal domain of Ire1 revealed an interface-1 (IF1) required for dimerization, and an interface-2 (IF2) providing a platform for the back-to-back association of dimers in higher oligomeric assemblies (Credle et al., 2005; Korennykh and Walter, 2012). Consistent with a previous report (van Anken et al., 2014), the formation of microscopically visible clusters of Ire1 is abolished by disrupting either IF1 or IF2 by mutation (T226A/F247A and W426A for IF1 and IF2, respectively) (Fig. 6A). Expectedly, lack of clustering correlates with an increased cellular sensitivity to DTT *(Suppl. Materials* Fig. S6).

**Figure 6.**
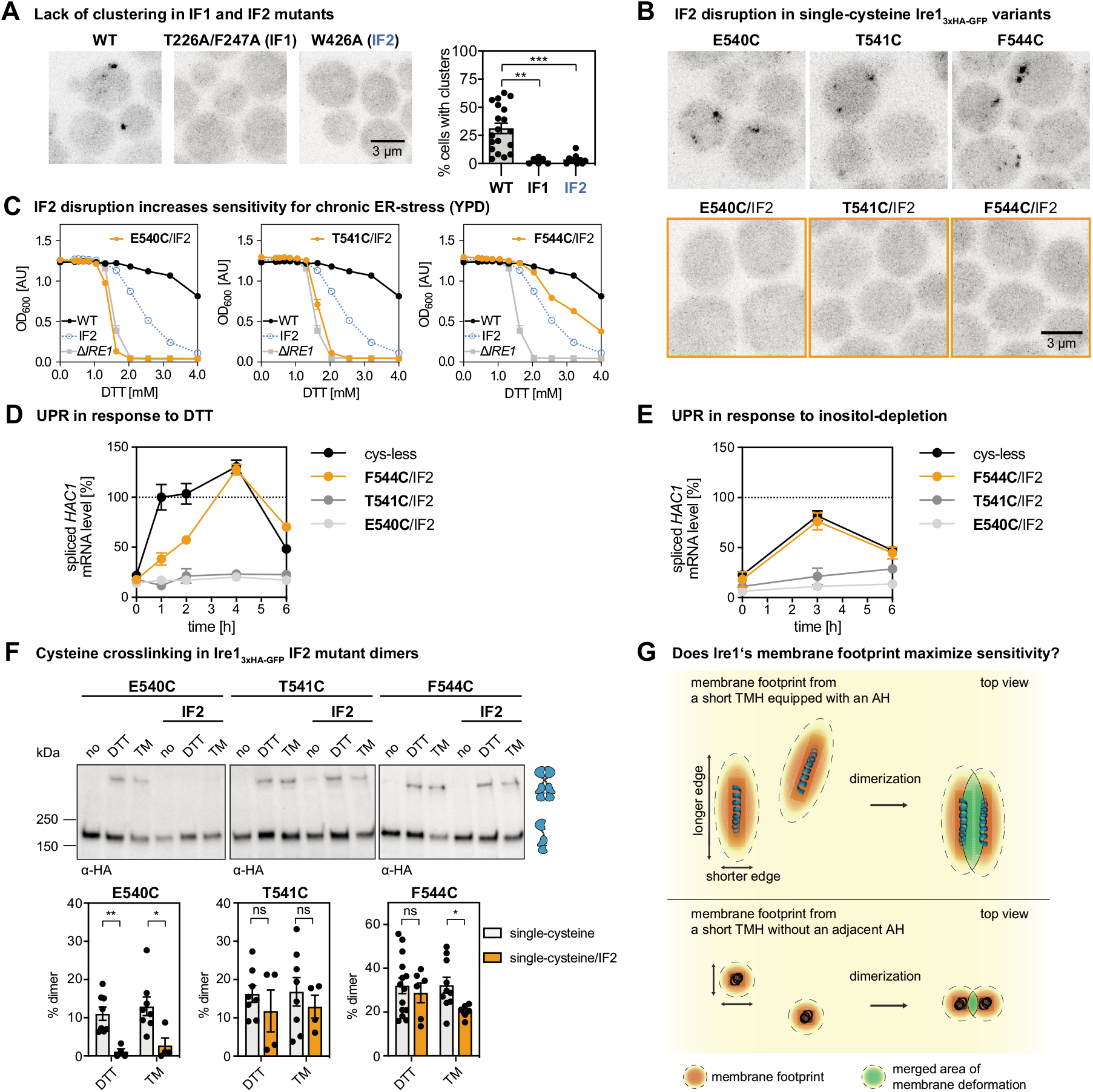
Crosslinking occurs within and across dimers of Ire1. (*A*) Indicated variants of an *IRE1* knock-in construct (Halbleib et al., 2017) were cultivated and stressed with 2 mM DTT for 1 h as described in Fig. 1D. A refined, automated counting of Ire1-containing clusters was performed as described in the *Suppl. Materials*. Data from Fig. 1D (WT: n = 13) were re-analyzed and pooled with new data (WT: n=6; T226A/F247A (IF1): n=6; W426A (IF2): n=10). All data are represented as the mean ± SEM. Significance was tested by an unpaired, two-tailed Student’s t test. ***p<0.001, **p<0.01. (*B*) Clustering in DTT-stressed cells was studied by confocal microscopy of the indicated single-cysteine variants with either an intact or disrupted IF2 (orange). Representative images from at least six independent fields of views are shown. For a quantitative analysis, see *Suppl. Materials* Fig. S6B. (*C*) ER-stress resistance of indicated cells was studied in rich medium containing different concentrations of DTT. Data for single-cysteine variants of Ire1 (E540C; T541C; F544C) all carrying an IF2-disrupting mutation (W426A) are plotted in orange. Reference data sets for cells lacking *IRE1* (gray) and cells producing either Ire1_3xHA-GFP_ WT or an IF2-disrupted variant are shown in black and blue, respectively. The data are from at least four independent experiments and represented as the mean ± SEM. (*D*) The level of the spliced *HAC1* mRNA was determined from the indicated strains by RT-qPCR after treating the cells with 2 mM DTT for the indicated times. The data are normalized to the level of the spliced *HAC1* mRNA in cysteine-less Ire1 after one hour of treatment. (*E*) The level of the spliced *HAC1* mRNA was determined for the indicated strains cultivated under inositol-depletion conditions. The data are normalized to the level of the spliced *HAC1* mRNA splicing caused by 2 mM DTT, as determined in (D). (*F*) The impact of the IF2-disrupting W426A mutation on crosslinking via the indicated single-cysteines was determined. Single- and double-mutant strains were subjected to the crosslinking procedure as in Fig. 3E. Data for the single mutant variants are re-plotted from Fig. 3F. All data are represented as the mean ± SEM derived from at least four independent experiments. Significance was tested by an unpaired, two-tailed Student’s t test. **<0.01, *p<0.05, ns: not significant. (*G*) Hypothetical model for Ire1’s exquisite sensitivity. The membrane-based oligomerization of Ire1 (blue) and unrelated single-pass membrane proteins (black) leads to the coalescence of deformed membrane regions (green). In the case of Ire1, a larger portion of the deformed membrane region can be shared upon dimerization, due to the ellipsoid membrane ‘footprint’ and an association via the longer edge of deformation (parallel to the major axis of the ellipse). According to this model, this maximizes the sensitivity of Ire1 to aberrant membrane properties when compared to unrelated single-pass membrane proteins, which can merge only a relatively small portion of their circular membrane ‘footprint’ upon dimerization.

By disrupting IF2 and leaving IF1 intact, we sought to uncover the contribution of dimeric and oligomeric assemblies to the crosslinking propensity in the TMH region. We focused on F544C marking the crossing-point of the X-shaped TMH region in dimeric Ire1, and on E540C and T541C in the vicinity. Upon disruption of IF2 (W426A), these single-cysteine variants failed to form microscopically visible clusters in stressed cells (Fig. 6B). The positioning of the engineered cysteine, however, had profound impact on the cellular resistance to DTT in rich medium. The F544C/IF2 double mutant rendered the respective cells more resistant than the IF2 mutant alone, while the T541C/IF2 and E540C/IF2 mutants were highly sensitive to DTT and indistinguishable from cells lacking *IRE1* altogether (Fig. 6C). Thus, the functional defect from the IF2 mutation can be alleviated or even aggravated by polar residues in the TMH region.

For interpreting these data, it is important to consider the timeframe of the different assays. Crosslinking is performed with microsomes isolated from acutely stressed cells, which were treated with either DTT or TM for only one hour. Similarity, clustering of Ire1 is studied by confocal microscopy in acutely cells stressed after one hour of treatment. The cellular resistance to DTT, however, is scored after 18 hours of cultivation. The acute proteotoxic stress caused by DTT or TM treatments has barely any impact on the cellular lipid composition under given conditions (Reinhard et al., 2020). Prolonged treatments, however, cause membrane aberrancies, which can dominate Ire1 activation (Promlek et al., 2011) and which are likely to affect the resulting ER stress resistance phenotype.

In order to further characterize the impact of the single-cysteine variants on Ire1 function, we determined the level of the spliced *HAC1* mRNA in time-course experiments with DTT-stressed cells (Fig. 6D). We find that the level of the spliced *HAC1* mRNA is upregulated in response to DTT-induced stress for cysteine-less Ire1 and F544C/IF2, but not for the E540C/IF2 and T541C/IF2 double mutants (Fig. 6D). Notably, we find that UPR activation is delayed for the F544C/IF2 double mutant compared to the cysteine-less control strain. Because membrane aberrancies caused by DTT manifest over a time course of several hours (Promlek et al., 2011), this suggests that the F544C/IF2 double mutant may respond predominantly to such membrane-based stresses. This interpretation is further confounded by the observation that the two other double mutants, E540C/IF2 and T541C/IF2 with mutations in the functionally critical AH (S526-V543) cannot respond to this type of prolonged DTT stress (Fig. 6D). In order cross-validate our interpretation, we studied the response of the same set of strains to lipid bilayer stress caused by inositol-depletion (Fig. 6E). While the F544C/IF2 mutation exhibited an almost identical response to inositol-depletion as the control strain, the two E540/IF2 and T541/IF2 variants showed a massively impaired response (Fig. 6E). Because E540 and T541 are part of the AH, these data underscore the central importance of the AH for sensing lipid bilayer stress. More importantly, however, these data suggest that membrane-sensitivity of Ire1 may be particularly important for dealing with prolonged forms of ER stress caused by proteotoxic agents.

Next, we subjected these double mutant variants to the crosslinking procedure (Fig. 6F). Crosslinking via E540C in DTT- and TM-stressed cells was abolished by the IF2 mutation, while the crosslinks observed for the T541C and F544C variant were only marginally affected (Fig. 6F). This suggests that the crosslinks of T541C and F544C are formed within Ire1 dimers, while E540C crosslinks across dimers. Importantly, these data validate not only the particular position of F544 at the crossing-point of two TMHs in two adjacent Ire1 protomers, they also provide direct, biochemical evidence that the unusual X-shaped transmembrane architecture might laterally associate to form higher-oligomers. Notably, such lateral ‘stacking’ of the transmembrane domain in signaling-active clusters would be consistent with the complex, elongated organization of clusters as recently observed by super-resolution microscopy for IRE1α (Belyy et al., 2020). On the functional level, our data show that the dimerization of Ire1 is not sufficient to mediate resistance to ER stress: the T541C/IF2 variant forms dimers that can be crosslinked (Fig. 6D), but it does not render cells more resistant to DTT than cells lacking Ire1 (Fig. 6C) nor does it upregulate the level of the *HAC1* mRNA in response to ER stress (Fig. 6D, E). A suitably positioned polar residue (here F544C) leaves the membrane-sensitive AH intact and increases the cellular ER stress resistance in the IF2/F544C double mutant compared to a single IF2 mutant (Fig. 6C). Thus, seemingly subtle chances in the TMH region can have substantial impact on the ER stress resistance phenotype especially, when the normal function of Ire1 is compromised.

## Discussion

Here, we establish a structural model of Ire1’s TMH region in signaling-active clusters (Fig. 4). In a previous study, we have established a model of Ire1’s monomeric TMH region (Halbleib et al., 2017), but its organization in dimers and higher oligomers, especially in the complex environment of the ER membrane, remained unexplored. Predicting a dimeric structure based on a model for the monomer is not trivial as the two protomers can be arranged in various ways and might undergo substantial conformational changes upon oligomerization. Based on a systematic cysteine crosslinking approach in native membranes and aided by MD simulations, we show that the neighboring TMHs in clusters of Ire1 organize in an X-shaped configuration.

Our model of the transmembrane organization provides intriguing insights into the membrane-deforming potential of Ire1 (Fig. 4A, B; *Suppl. Materials* Fig. S4, Movie S1). Positively charged residues at the cytosolic end of the TMH (Fig. 3A) and the previously identified ER-luminal AH (Halbleib et al., 2017) cooperate in squeezing the lipid bilayer (Fig. 4A, B; *Suppl. Materials* Fig. S4A). This deformation is most prominent at the intersection of the two protomers reaching almost to the level of the lipid bilayer center (Fig. 4B, *Suppl. Materials* Movie SI). Membrane squeezing and the associated disordering of lipid acyl chains come at energetic costs, which are affected by the composition and collective physicochemical properties of the surrounding bilayer (Radanović et al., 2018; Covino et al., 2018). The higher this cost (e.g. due to increased lipid saturation, inositol-depletion, or membrane aberrancies from prolonged proteotoxic stresses), the higher the free energy gain from coalescing these regions and thus the propensity of Ire1 to oligomerize.

The specific way each membrane protein locally deforms the bilayer, referred to as membrane ‘footprints’ (Haselwandter and Mackinnon, 2018) or ‘fingerprints’ (Corradi et al., 2018), could be at the origin of membrane-sensitivity and, more generally, control the organization of supramolecular assemblies (Corradi et al., 2018). Is it possible that the unusual TMH region of Ire1 and its resulting footprint serves a specific function? We speculate that the combination of a short TMH with an AH inserting deep into the bilayer contributes to Ire1’s exquisite sensitivity to aberrant ER membrane stiffening. The region of membrane compression around monomeric Ire1 is, when viewed form the top, not of circular shape but ellipsoid due to the membrane-inserted AH (Fig. 6G) (Halbleib et al., 2017). Based on simple geometric considerations, it is conceivable that the total extent of membrane deformation contributing to the free energy of dimerization depends on how precisely the two TMH regions are arranged towards each other. Our structural model of the dimeric TMH suggests that the two protomers associate via the longer edge of membrane deformation (parallel to the major axis of the ellipse) (Fig. 4A, B) thereby maximizing the area of coalescence (Fig. 6G, top) and minimizing the free energy. We speculate that Ire1 is more responsive to aberrant membrane stiffening than other single-pass transmembrane proteins with short TMHs but without AHs. Because these proteins also lack the characteristic ellipsoid shape of membrane deformation (Kaiser et al., 2011), they coalesce only a smaller area of their footprints upon dimerization (Fig. 6G, bottom). It will be intriguing to study the membrane-driven dimerization and oligomerization of Ire1 side-by-side with other single-pass membrane proteins exhibiting distinct membrane footprints using advanced microscopic tools such as single-molecues photobleaching (Chadda et al., 2016).

Our data also provide evidence that crosslinking can occur across dimers of Ire1 (Fig. 6F), thereby suggesting that the X-shaped dimeric arrangements of the TMH region can laterally associate and ‘stack’ in the plane of the membrane. We propose that it is the characteristic, ellipsoid shape of membrane deformation by monomeric Ire1 and the unusual mode of dimerization and oligomerization, which maximizes the sensitivity of Ire1 to aberrant membrane properties.

Our structural and functional analyses suggest that the oligomeric state of Ire1 is stabilized by the overall transmembrane architecture and the membrane-embedded AH, but not by specific interactions between residues in the TMH. Disrupting the AH, which also disrupts transmembrane architecture (Covino et al., 2018), increases the cellular sensitivity to ER stress (Figure 5C). In contrast, the F544A mutation at the intersection of neighboring TMHs causes no functional defect (Fig. 5A,B). Instead of maximizing the interface between the TMHs for forming a more stable protein:protein interaction, they are kept in a configuration where only a few TMH residues can contact the opposing protomer. However, they are driven together via a membrane-based mechanism and thus particularly sensitive to the properties of the surrounding membrane (Covino et al., 2018).

Strikingly, our data provide evidence that different forms of ER stress converge in a single, overall transmembrane architecture of Ire1. We observed remarkably similar crosslinking patterns in the context of lipid bilayer stress and proteotoxic stress (Fig. 3F). This suggests that the X-shaped configuration in the TMH region is maintained in the signaling-active clusters even under largely distinct conditions of ER stress. Neither the oligomerization of Ire1 *per se* nor lipid bilayer stress seem to cause major conformational changes in the TMH region of the individual protomers. Based on our data, we speculate that Ire1 mounts a single response to different types of ER stress, however, with distinct temporal patterns of activation. Proteotoxic stress caused by DTT or TM is characterized by two phases: An early phase of a rapid UPR activation with little to know changes in the lipid composition and a second, slower phase characterized by a build-up of membrane-aberrancies (Promlek et al., 2011; Reinhard et al., 2020). While these membrane aberrancies remain poorly characterized, they serve as a robust signal for Ire1 activation (Fig. 6D) (Promlek et al., 2011). The lipid bilayer stress caused from inositol-depletion, in contrast, lacks the early phase of UPR activation. It manifests slowly and causes a distinct temporal pattern of UPR activation (Fig. 6D, E). It will be interesting to study, if different temporal patterns of UPR activation are sufficient to give rise to largely distinct transcriptional programs or if -alternatively-Ire1 can custom-tailor its output via yet unknown mechanisms (Hetz et al., 2020; Ho et al., 2020; Fun and Thibault, 2020).

Our crosslinking data suggest a similar transmembrane architecture in Ire1 in response to proteotoxic and and lipid bilayer stress (Gardner and Walter, 2011; Halbleib et al., 2017). While we cannot formally exclude conformational changes in other parts of the protein, we do not find evidence that Ire1 custom-tailors its signaling-output via conformational changes in the TMH region. Based on our crosslinking data and the observed temporal patterns of activation for different mutants of Ire1 (Fig. 6E), we suggest that the complex metabolic, transcriptional, and non-transcriptional adaptations to different forms of ER-stress do not reflect distinct functional modes of Ire1. Instead, we propose that different degrees of oligomerization and different rates of Ire1 activation and inactivation are sufficient to drive differently stressed cells into distinct physiological states.

Our combined results lead to the following model of UPR activation. Both accumulating unfolded proteins and lipid bilayer stress lead to the oligomerization of Ire1 and the formation of signaling-active clusters (Korennykh and Walter, 2012). Under these conditions, the cytosolic effector domains ‘follow’ the oligomerization of the ER-luminal domain and the TMH region. A large diversity of ER-luminal and cytosolic interactors including chaperones can tune and specify the activity of mammalian UPR transducers (Sepulveda et al., 2018; Amin-Wetzel et al., 2017). This may reflect a way to custom-tailor the globally acting UPR to different cell types with distinct protein folding requirements at steady state and during differentiation. Lipid bilayer stress activates the UPR in both yeast and mammals via a membrane-based mechanism and does not require the binding of unfolded proteins to the ER-luminal domain and/or associated chaperones (Promlek et al., 2011; Halbleib et al., 2017; Volmer et al., 2013). Furthermore, our findings underscore the importance of Ire1’s membrane-sensitivity to deal with the stress caused by prolonged cellular treatments with proteotoxic agents (Promlek et al., 2011). Our data from direct, crosslinking experiments suggest that both proteotoxic and lipid bilayer stress converge in a single overall architecture of the TMH region. We propose that Ire1’s distinct signaling outputs to different forms of ER stress reflect a different temporal pattern of Ire1 activation rather than different qualities of signaling.

## Materials and Methods

### Reagents, Antibodies, Strains, and Plasmids

All chemicals and reagents used in this study were purchased from Sigma Aldrich, Carl Roth or Millipore and are of analytical or higher grade. The following antibodies were used: mouse anti-Flag monoclonal (M2) (Santa Cruz), rat anti-HA monoclonal (3F19) (Roche), mouse anti-Dpm1 monoclonal (5C5A7) (Life Technologies), mouse anti-Pgk1 (22C5D8) (Life Technologies), mouse anti-MBP monoclonal (NEB), anti-mouse-HRP (Dianova), anti-rat-HRP (Dianova). All strains and plasmids used in this study are listed in *Suppl. Materials* Table S1, S2).

### Generation of a cysteine-less construct and a Flag-tag variant of *IRE1*

The construction of a cysteine-less construct of *IRE1* is based on a previously described knock-in construct (Halbleib et al., 2017). This construct comprises the *IRE1* promotor (−1 to −551 bp), the *IRE1* gene including a coding sequence for a 3xHA tag and a monomeric version of yeGFP (A206R^yeGFP^) inserted at the position of H875, and the *IRE1* endogenous 5′ terminator on the plasmid pcDNA3.1-IRE1-3xHA-GFP (Halbleib et al., 2017). A cysteine-less variant was generated by site-directed mutagenesis. Cysteine 48 (C48^yeGFP^) of the monomeric yeGFP was substituted to serine, while cysteine 70 (C70^yeGFP^) remained in the final construct (Costantini et al., 2015; Ormö et al., 1996). Single-cysteine variants were generated by site-directed mutagenesis.

Plasmids encoding either single-cysteine variants or cysteine-less Ire1 (Table S2) were linearized using *Hind*III and *Xho*I restriction enzymes and used for transforming our previously established cloning strain lacking both the *IRE1* gene and its promotor. Strains used in this study are listed in Table S1. Additionally, a Flag-tagged cysteine-less Ire1 version based on the *CEN*-based Ire1 construct from the pPW1628/pEv200 plasmid was generated. The 3xHA epitope tag in the knock-in construct was replaced by a 3xFlag epitope tag using the Q5 site-directed mutagenesis kit (NEB). The newly generated knock in sequence was amplified in a multi-step PCR reaction adding the terminator sequence from the pEv200 plasmid and *BssH*I and *Hind*III restriction site. The transfer of the *IRE1_3xFlag-GFP_* sequence in the *CEN*-based pPW1628/pEv200 plasmid was performed using *BssH*I/*Hind*III restriction sites.

### Cultivation and live cell confocal microscopy

The yeast strains were cultivated at 30°C on agar plates containing SCD complete medium or selection medium. Liquid yeast cultures either in SCD or YPD (the pH of the medium was not adjusted) were inoculated with a single colony and typically cultivated at 30°C for a minimum of 18 h to reach the stationary phase. This overnight culture was used to inoculate a fresh culture to an OD_600_ = 0.2, which was cultivated until the mid-exponential phase. For microsomal membrane preparation, stationary cells were used to inoculate a fresh culture in SCD complete medium to an OD_600_ of 0.2. After cultivation at 30°C to an OD_600_ of 0.7, the cells were either left untreated or stressed with either 2 mM DTT or 1.5 μg/ml Tunicamycin for 1 h. For inositol depletion, exponentially growing cells were washed with SCD complete w/o inositol and then used to inoculate the main culture to an OD_600_ of 0.5 in SCD complete w/o inositol, which was further cultivated for 3 h.

### Live cell confocal microscopy and image analysis

A fresh culture in SCD medium was inoculated to an OD_600_ = 0.2 and cultivated for 5 to 5.5 h at 30°C and under constant agitation at 220 rpm. To induce ER-stress, DTT was added to a final concentration of 2 mM followed by additional cultivation for 1 h. The cells were harvested by centrifugation and mounted on microscopic slides coated with a thin layer of SCD containing 1.5% agarose for immobilization. Microscopy was performed using a Zeiss LSM 780 confocal laser scanning microscope (Carl Zeiss AG) with spectral detection and a Plan-Apochromat 63x 1.40 NA oil immersion objective. GFP fluorescence was excited at 488 nm and the emission was detected between 493 and 598 nm. Transmission images were simultaneously recorded using differential interference contrast (DIC) optics. Z-stacks (450 nm step-size, 62.1 μm pinhole size) were recorded. When multiple fluorophores were imaged (*Suppl. Materials* Fig S6F), GFP was excited at 488 nm, dsRed at 561 nm and emission was detected at 493-557 nm and 592-704 nm respectively. For multi-fluorophor images a Z-stack step-size of 372 nm with a pinhole diameter of 80.3 μm was used. Image stacks were corrected for potential xy-drift using the Fiji plugin StackReg (Thévenaz et al., 1998; Schindelin et al., 2012). Maximum intensity and sum projections were created, while the contrast was adjusted equally for all images using Fiji (Schindelin et al., 2012). Individual cells and clusters of Ire1 were identified by automated segmentation using CellProfiler (McQuin et al., 2018). In brief, the cellular areas were determined for each image based on sum projections of recorded z-stacks and the cellular autofluorescence. After smoothing with a median filter, potential cells were identified by global thresholding (minimum cross entropy). Objects outside the diameter restraint of 1.9 - 6.3 μm were discarded. Cells being too bright (a high autofluorescence indicates cell death) were omitted from further analysis if the mean intensity of a potential cell exceeded the mean intensity of all potential cells within an image by more than 30%. Clusters of Ire1 within cells were identified in maximum intensity projections using a threshold of 1.5 times the mean intensity of the identified cells. Potential clusters outside the diameter range 0.3 - 0.9 μm were discarded. The strain RE773 *IRE1*-3xHA-yeGFP W426A E540C *single cysteine* showed substantial signs of cell death (increased autofluorescence) when challenged with DTT. Therefore, all microscopic images represented in and used for Fig. 6 and *SI Appendix* Fig. S6B were reanalyzed and subjected to more stringent parameters to avoid false positive identifications of Ire1 clusters. Cells were not considered, if their mean intensity was 10% above average. Structures with diameters from 0.3 - 1.2 μm, were initially allowed as potential clusters, but only counted if their maximum intensity was at least 2.5-times higher than the mean intensity of the respective cell. Furthermore, if more than 3.5% of a cell area was covered by potential clusters, the cell was considered as unfit and counted as free of clusters.

### Assaying the resistance to ER-stress

The cellular resistance to ER-stress caused by DTT was assayed using a sensitive growth assay (Halbleib et al., 2017). Stationary overnight cultures were used to inoculate a fresh culture to an OD_600_ of 0.2. After cultivation for 5 to 7 h at 30°C the cells were diluted with pre-warmed medium to an OD_600_ of 0.05. 50 μl of these diluted cultures were mixed in a 96-well plate with 180 μl of medium and 20 μl of a DTT dilution series leading to a final concentration of DTT between 0 and 2 mM and 0 and 4 mM, respectively. After incubation at 30°C for 18 h, the cultures were thoroughly mixed and 200 μl of the cell suspension were transferred to a fresh 96-well plate for determining the density of the culture via spectrophotometers using the OD_600_/OD_620_.

### RNA preparation, cDNA synthesis and quantitative real-time (qPCR) PCR analysis

The level of the spliced *HAC1* mRNA and the *PDI1* mRNA in stressed and unstressed cells was determined via RT-qPCR using Oligo(dT) primers, the Superscript™ II RT protocol (Invitrogen), the ORA qPCR Green ROX L Mix (HighQu) and a Piko Real PCR system (Thermo Scientific). The RNA was prepared from 5 OD equivalents of stressed and unstressed cells using the RNeasy Plus RNA Isolation Kit (Qiagen). 500 ng RNA of the total isolated RNA were used as a template for the synthesis of cDNA using Oligo(dT) primers and the Superscript™ II RT protocol (Invitrogen). qPCR was performed using ORA qPCR Green ROX L Mix (HighQu) in a Piko Real PCR system (Thermo Scientific). The following primers were used at a final concentration of 400 nM:

*HAC1s* forward primer: 5’ – CTTTGTCGCCCAAGAGTATGCG – 3’
*HAC1s* reverse primer: 5’ – ACTGCGCTTCTGGATTACGC – 3’
*ACT1* forward primer: 5’ – TGTCACCAACTGGGACGATA – 3’
*ACT1* reverse primer: 5’ – AACCAGCGTAAATTGGAACG – 3’
*PDI1* forward primer: 5’ – GATCGATTACGAGGGACCTAGA – 3’
*PDI1* reverse primer: 5’ – GCGGAGGGCAAGTAAATAGAA – 3’

The qPCR program included the following steps: 1) 95°C, 15 min; 2) 95°C, 20 sec; 3) 58°C, 20 sec; 4) 72°C, 30 sec; 5) 72°C, 5 min; steps 2-4 were repeated 40 times. For quantifying the level of the *PDI1* mRNA and the spliced *HAC1* mRNA, we used the comparative ΔΔCT method using normalization to *ACT1* levels (StepOnePlus™ user Manual, Applied Biosystems).

For amplifying both cDNAs generated from the spliced and unspliced *HAC1* mRNA, we used the following primer at a final concentration of 400 nM and previously established polymerase chain reaction (PCR) conditions (Promlek et al., 2011).

*HAC1* splicing forward primer: 5’-TACAGGGATTTCCAGAGCACG-3’
*HAC1* splicing reverse primer: 5’-TGAAGTGATGAAGAAATCATTCAATTC-3‘

### Preparation of cell lysates and immunoblotting

Lysates were prepared from exponentially growing cells, which were harvested by centrifugation (3.000xg, 5 min, 4°C) and then washed once with ddH_2_O and once with PBS. During washing, the cells were transferred into 1.5 ml reaction tubes allowing for a more rapid centrifugation (8.000xg, 20 sec, 4°C). The tubes with the washed cell pellet were placed in a −80°C freezer and stored until further use. For preparing the lysate, either 5 or 20 OD equivalents were resuspended in 400 μl or 1000 μl lysis buffer (PBS containing 10 μg/ml chymostatin, 10 μg/ml antipain, 10 μg/ml pepstatin), respectively. After addition of either 100 μl or 500 μl of zirconia beads, respectively, the cells were disrupted by bead beating for 5 min at 4°C. Four volumes units of the resulting lysate were mixed with one volume of 5x reducing sample buffer (8 M urea, 0.1 M Tris-HCl pH 6.8, 5 mM EDTA, 3.2% (w/v) SDS, 0.15% (w/v) bromphenol blue, 4% (v/v) glycerol, 4% (v/v) β-mercaptoethanol) and then incubated at 95°C for 10 min for fully unfolding and solubilizing the proteins therein. 0.1 OD equivalents of the resulting sample was subjected to SDS-PAGE and the proteins were separated on 4-15% Mini-PROTEAN-TGX strain-free gels (BioRad). For subsequent immuno-blotting, proteins were transferred from the gel to methanol-activated PVDF membranes using semi-dry Western-Blotting. Specific proteins were detected using antigen-specific primary antibodies, HRP-coupled secondary antibodies, and chemiluminescence. The percentage of crosslinked dimer was determined via densitometry with Fiji (Schindelin et al., 2012) using the bands corresponding to the monomeric and covalently-crosslinked protein.

### Microsomal membrane preparation

80 OD_600_ equivalents were harvested from a mid-exponential culture by centrifugation (3.000xg, 5 min, 4°C), washed with PBS, and stored at −80°C. All steps of membrane fractionation were performed on ice or at 4°C. Cells were resuspended in 1.5 ml lysis buffer (50 mM HEPES pH 7.0, 150 mM NaCl, 1 mM EDTA, 10 μg/ml chymostatin, 10 μg/ml antipain, 10 μg/ml pepstatin). For cysteine crosslinking experiments, a buffer without EDTA was used. After cell disruption using zirconia beads (Roth) and a bead beater (2 × 5 min), cell debris was removed by centrifugation (800x g, 5 min, 4°C) and (5,000 x g, 10 min, 4°C). The supernatant was centrifuged (100.000x g, 45 min, 4°C) to obtain crude microsomes in the pellet. Microsomes were resuspended in 1.4 ml lysis buffer, sonicated for homogenization (50%, 5×1sec, MS72 tip on a sonifier cell disrupter from Branson Ultrasonic), snap frozen in liquid N_2_, and stored in aliquots at −80°C.

### Test of membrane integration

The cleared supernatant of a 5.000xg step was divided into equal parts, which were then mixed with an equal volume of lysis buffer supplemented with 0.2 M Na_2_CO_3_ resulting in a final pH of 11, 5 M urea, 2% Triton X-100 or without additional additives. After incubation for 1 h on a rotator, these samples were centrifuged (100,000x g, 45 min, 4°C) to separate soluble from insoluble material. The supernatant and pellets from these fractions corresponding to 0.2 OD equivalents were further analyzed by SDS-PAGE and immunoblotting.

### CuSO_4_-induced cysteine crosslinking

Microsomes were thawed on ice. 8 μl microsomes (1 ± 0.2 mg/ml protein) were mixed either with 2 μl of 50 mM CuSO_4_ or 2 μl ddH_2_O and then incubated for 5 min on ice. The reaction was stopped with 8 μl of membrane sample buffer (4 M urea, 50 mM Tris-HCl pH 6.8, 1.6 % (w/v) SDS, 0.01% (w/v) bromophenol blue, 2% (v/v) glycerol) containing 125 mM EDTA and 250 mM NEM. The samples were analyzed by SDS-PAGE and immunoblotting. See the *Suppl. Materials* for further details.

### Immunoprecipitation from microsomes after CuSO_4_-induced cysteine crosslinking

300 μl of microsomes with a typical protein concentration of 1 mg/ml were incubated with 12.5 μl 250 mM CuSO_4_ (final concentration of 10 mM) for 5 min on ice. The reaction was stopped by adjusting the sample to a final concentration of 50 mM EDTA and 111 mM NEM by adding 30 μl of 0.5 M EDTA stock solution and 44 μl of 1 M NEM stock solution, respectively. The final volume was adjusted to 1.3 ml with lysis buffer with a final concentration of 5 mM EDTA. The CuSO_4_ concentration was thus reduced to 2.4 mM and the NEM concentration to 33.6 mM, respectively. After crosslinking, the microsomes were solubilized using 2% Triton X-100 and incubated for 1 h at 4°C under constant agitation. Insoluble material was removed by centrifugation (20.000x g, 10 min, 4°C). The resulting supernatant was incubated with 8 μl Flag beads (Sigma Aldrich), equilibrated with IP wash buffer (lysis buffer + 5 mM EDTA + 0.2 % Triton X-100), for 3 h under constant shaking. Flag beads were washed five times with IP wash buffer by centrifugation (8.000xg, 30 sec, 4°C). For elution, the Flag beads were incubated with 10 μl IP-Wash and 10 μl 5x reducing sample buffer for 5 min at 95°C, which did not disrupt the disulfide bond formed between to protomers of Ire1. These samples were analyzed by SDS-PAGE and immunoblotting.

### Modelling of the transmembrane dimer of Ire1 and MD simulations

The dimeric TMH region of Ire1 was modeled using a 56 amino-acid long peptide 516-SRELD EKNQNSLLLK FGSLVYRIIE TGVFLLLFLI FCAILQRFKI LPPLYVLLSK I-571. We extracted an equilibrated, monomeric configuration of the peptide from a previously performed 10 μs long equilibrium MD simulation. We duplicated the configuration in order to create a new system containing two identical protomers. We then rotated and translated one of the two protomers to form a dimer structure, such that the two F544 faced each other with the distance between their Cβ atoms at around 0.7 nm. A short energy minimization in solution resolved all steric clashes between sidechains. The structure of the model dimer was prepared by using gromacs/2019.3 tools (Abraham et al., 2015) and VMD (Humphrey et al., 1996). We used Charmm-GUI (Wu et al., 2014; Lee et al., 2016) to reconstitute the dimer in a bilayer containing 248 POPC and 62 cholesterol molecules modelled in the Charmm36m force-field (Klauda et al., 2010; Best et al., 2012). We solvated the system with 24813 TIP3P water molecules, 72 chloride and 66 sodium ions, corresponding to a salt concentration of 150 mM.

### Equilibrium and restrained simulations of the dimer model

After an initial energy minimization and quick relaxation, we equilibrated the dimer model in the bilayer. We first ran a 50 ns long simulation restraining the position of protein atoms by using harmonic potentials with force-constants (in units of kJ mol^−1^ nm^−2^) of 500 for backbone atoms and 200 for side-chain atoms. We then ran further 50 ns lowering the force-constants to 200 and 50, respectively. After this equilibration, we relieved all restraints and ran a 1000 ns long MD simulation, where the system evolved according to its unbiased dynamics. We ran both the restrained equilibration and unbiased production simulation in gromacs/2019.3 using a time step of 2 fs. Electrostatic interactions were evaluated with the Particle-Mesh-Ewald method (Essmann et al., 1995). We maintained a constant temperature of 303 K (Bussi et al., 2007), applying separate thermostats on the protein, membrane, and solvent with a characteristic time of 1 ps. We applied the semi-isotropic Berendsden barostat (Berendsen et al., 1984) for the restrained equilibration, and the Parrinello-Rahman barostat (Parrinello and Rahman, 1981) for the production runs, acting separately on the x-y plane and z direction to maintain a constant pressure of 1 atm, and with a characteristic time of 5 ps. We constrained all hydrogen bonds with the LINCS algorithm (Hess et al., 1998). Molecular visualizations were obtained with VMD and rendered with Tachyon.

### Data representation and replicates

All data are represented as the average ± SEM if not stated otherwise. The number of the biological and technical replicates are provided in the *Suppl. Materials*. Statistical tests were performed with Prism 8 for macOS Version 8.4.0.

## Data availability statement

All data discussed in the paper are included in this published article and in the *Suppl. Materials*. Additional materials, such as qPCR data, microscopy data, and the immunoblots contributing to the bar diagrams in Fig. 3F, 5B, 5D, and 6D as well as in *Suppl. Materials* Fig. S5B have been deposited to Mendeley Data (DOI:10.17632/s52vt8spmc.1).

## Acknowledgments

This work was supported by the Deutsche Forschungsgemeinschaft (SFB807 ‘Transport and Communication across Biological Membranes’ to R.E. and G.H.; SFB894 ‘Ca^2+^-Signals: Molecular Mechanisms and Integrative Functions’ to R.E). RE was supported by the Volkswagen Foundation (Life?, #93089). This project has received funding from the European Research Council (ERC) under the European Union’s Horizon 2020 research and innovation program (grant agreement No. 866011). R.C. and G.H. were supported by the Max Planck Society. We thank Kristina Halbleib for her important contributions during the early phase of the project and David Ron for critically reading the manuscript and helpful discussions. We thank Sebastian Schuck and Dimitrios Papagiannidis for providing the pSS455 plasmid.

## Supplementary Material

**Number of independent experiments for each dataset**

**Fig. 1B:** *Left panel*: Δ*IRE1*, WT: n=20 (data from four individual colonies with technical replicates); cysteine-less: n=12 (technical replicates from four individual colonies). *right panel*: Δ*IRE1*: n=14 (data from three individual colonies with technical replicates); cysteine-less: n=12 (technical triplicates from four individual colonies); WT: n=9 (technical triplicates from three individual colonies).

**Fig. 1C:** *Left panel*: WT -DTT: n= 4; WT +DTT: n=6; cysteine-less -DTT: n=6; cysteine-less +DTT: n=5.

Right panel: WT -TM: n=4; WT +TM: n=5; cysteine-less -TM: n=6; cysteine-less +TM: n=4.

**Fig. 1D:** WT -DTT: n=6 (fields of view, total number of cells = 172), WT +DTT: n=13 (fields of view, total number of cells = 302), cysteine-less -DTT: n=6 (fields of view, total number of cells = 209), cysteine-less +DTT: n=12 (fields of view, total number of cells = 326).

**Fig. 3C: Ire1 variants +DTT**(biological replicates measured in technical duplicates); WT +DTT: n=5; cysteine-less +DTT: n=6; E540C +DTT: n=6; T541C +DTT: n=3; G542C +DTT: n=6; V543C +DTT: n=9; F544C +DTT: n=9; L545C +DTT: n=3; L546C +DTT: n=9; L547C +DTT: n=3; F548C +DTT; n=3; L549C +DTT: n=6; I550C +DTT: n=3; F551C +DTT: n=5; C552 +DTT: n=3.

**Ire1 variants -DTT**(biological replicates measured in technical duplicates) (WT-DTT: n=6; WT +DTT: n=5) as in Fig. 3C; cysteine-less -DTT: n=5; E540C -DTT: n=6; T541C -DTT: n=3; G542C -DTT: n=6; V543C -DTT: n=9; F544C -DTT: n=9; L545C -DTT: n=3; L546C -DTT: n=9; L547C -DTT: n=3; F548C -DTT; n=3; L549C -DTT: n=6; I550C -DTT: n=3; F551C -DTT: n=6; C552 -DTT: n=3.

**Fig. 3D: Inositol-depleted cells**(biological replicates measured in technical duplicates) (normalized to WT + DTT: n=5 as in Fig. 3C); WT -INO: n=3; cysteine-less -INO: n=3; E540C –INO: n=3; G542C –INO: n=3; F544C -INO: n=3; L546C -INO: n=3; L547C -INO: n=3; L549C –INO: n=3.

**Unstressed cells**(biological replicates measured in technical duplicates) (normalized to WT + DTT: n=5 as in Fig. 3C); WT SCD: n=3; cysteine-less SCD: n=3; E540C SCD: n=3; G542C SCD: n=3; F544C SCD: n=3; L546C SCD: n=3; L547C SCD: n=3; L549C SCD: n=3.

**Fig. 3F: DTT-stressed cells** E540C, T541C: n=8 (data from four individual colonies with technical replicates); G542C, V543C, L545C, L546C, L547C, F548C, L549C, I550C, F551C: n=4 (technical duplicates from two individual colonies); F544C: n=14 (technical duplicates from seven individual colonies); C552: n=11 (technical duplicates from six individual colonies).

**TM-stressed cells** E540C, T541C: n=8 (data from four individual colonies with technical replicates); G542C, V543C, L545C, L546C, L547C, F548C, L549C, I550C, F551C: n=4 (technical duplicates from two individual colonies); F544C: n=10 (data from five individual colonies with technical replicates); C552: n=8 (data from four individual colonies with technical replicates).

**Inositol depletion** E540C-L545C, F548C, F551C, C552: n=6 (data from three individual colonies with technical replicates); L546: n=5 (data from three individual colonies with technical replicates); L547C, L549C, I550C: N=4 (data from two individual colonies with technical replicates).

**Fig. 5A:**Δ*IRE1*, cysteine-less: n=12 (data from two individual colonies with technical replicates), F544A C552: n=12 (data from four individual colonies with technical replicates).

**Fig. 5B: DTT-stressed cells** C552: n=11 (data from six individual colonies with technical replicates – identical with data in Fig. 3F), F544A C552: n=4 (data from two individual colonies with technical replicates).

**TM-stressed cells** C552: n=8 (data from four individual colonies with technical replicates – identical with data in Fig. 3F), F544A C552: n=4 (data from two individual

**Fig. 5C:**Δ*IRE1*, cysteine-less: n=12 (technical replicates from two individual colonies, identical with data in Fig. 5A), F531R C552: n=9 (technical triplicates from three individual colonies).

**Fig. 5D: DTT-stressed cells** C552: n=11 (data from six individual colonies with technical replicates – identical with data in Fig. 3F), F531R C552: n=4 (data from two individual colonies with technical replicates).

**TM-stressed cells** C552: n=8 (data from four individual colonies with technical replicates – identical with data in Fig. 3F), F531R C552: n=4 (data from two individual colonies with technical replicates).

**Fig. 6A:** *reused for analysis*: WT: n=13 (fields of view, identical with data in Fig. 1D) with the total number of cells = 293; *additional data*: WT: n=6 (fields of view) with the total number of cells = 148). T226A/F247A (IF1): n=6 (fields of view, total number of cells = 154), W426A (IF2): n=10 (fields of view, total number of cells = 329).

**Fig. 6C:** *Plotted in all three panels:* WT: n=6 (technical replicates from two individual colonies); IF2: n=12 (technical replicates from two individual colonies); Δ*IRE1*: n=6 (technical replicates).

*Left panel*: E540C/IF2: n=4 (technical replicates).

*Middle panel*: T541C/IF2: n=5 (technical replicates).

*Right panel*: F544C/IF2: n=6 (technical replicates).

**Fig. 6D: DTT-stressed cells** cysteine-less, E540C/IF2, T541C/IF2, F544C/IF2: n=3 (data from three individual colonies, each measured in technical duplicates) for each time point (0 h, 1 h, 2 h, 4 h, 6 h).

**Fig. 6E: Inositol-depleted cells** cysteine-less, E540C/IF2, T541C/IF2, F544C/IF2: n=3 (data from three individual colonies, each measured in technical duplicates) for each time point (0 h, 3 h, 6 h).

**Fig. 6F: DTT-stressed cells** E540C, T541C: n=8 (data from four individual colonies with technical replicates, identical with data in Fig. 3F), F544C: n=14 (technical duplicates from seven individual colonies, identical with data in Fig. 3F); E540C/IF2, T541C/IF2: n=4 (data from two individual colonies with technical replicates), F544C/IF2: n=6 (data from three individual colonies with technical replicates).

**TM-stressed cells** E540C, T541C: n=8 (data from four individual colonies with technical replicates, identical with data in Fig. 3F), F544C: n=10 (data from five individual colonies with technical replicates, identical with data in Fig. 3F); E540C/IF2, T541C/IF2: n=4 (data from two individual colonies with technical replicates), F544C/IF2: n=6 (data from three individual colonies with technical replicates).

**Fig. S1E**: *Left panel*: WT -DTT: n= 5; WT +DTT: n=5; cysteine-less -DTT: n=6; cysteine-less +DTT: n=6.

*Right panel*: WT -TM: n=5; WT +TM: n=5; cysteine-less -TM: n=6; cysteine-less +TM: n=3.

**Fig. S3A: ER-stress resistance / growth assay**

OD_620_: Δ*IRE1*: n=20 (technical triplicates from three individual colonies, identical with data in Fig. 1B left panel);

cysteine-less: n=12 (technical triplicates from four individual colonies, identical with data in Fig. 1B left panel);

E540C – L546C: n=6 (technical triplicates from two individual colonies);

C552: n=12 (technical triplicates from four individual colonies);

OD_600_: Δ*IRE1*: n=10 (technical replicates from three individual colonies);

cysteine-less: n=22 (technical triplicates from four individual colonies);

L547C, L549C, I550C,F551: n=6 (technical triplicates from two individual colonies);

F548C: n=5 (technical replicates from two individual colonies).

**Fig. S3F: DTT-stressed cells:**

WT: n=13 (fields of view, total number of cells = 302) (as in Fig. 1D),

cysteine-less: n=12 (fields of view, total number of cells = 326) (as in Fig. 1D),

E540C: n=15 (fields of view, total number of cells = 359),

T541C: n=10 (fields of view, total number of cells = 206),

G542C: n=6 (fields of view, total number of cells = 124),

V543C: n=8 (fields of view, total number of cells = 223),

F544C: n=19 (fields of view, total number of cells = 439),

L545C: n=12 (fields of view, total number of cells = 181),

L546C: n=14 (fields of view, total number of cells = 399),

L547C: n=8 (fields of view, total number of cells = 203),

F548C: n=10 (fields of view, total number of cells = 279),

L549C: n=9 (fields of view, total number of cells = 212),

I550C: n=8 (fields of view, total number of cells = 232),

F551C: n=5 (fields of view, total number of cells = 152),

C552: n=7 (fields of view, total number of cells = 188).

**unstressed cells:**

WT: n=6 (fields of view, total number of cells = 172) (as in Fig. 1D),

cysteine-less: n=6 (fields of view, total number of cells = 209) (as in Fig. 1D),

E540C: n=4 (fields of view, total number of cells = 130),

T541C: n=7 (fields of view, total number of cells = 153),

G542C: n=7 (fields of view, total number of cells = 188),

V543C: n=4 (fields of view, total number of cells = 121),

F544C: n=4 (fields of view, total number of cells = 108),

L545C: n=5 (fields of view, total number of cells = 101),

L546C: n=3 (fields of view, total number of cells = 111,

L547C: n=4 (fields of view, total number of cells = 106),

F548C: n=5 (fields of view, total number of cells = 143),

L549C: n=4 (fields of view, total number of cells = 103),

I550C: n=5 (fields of view, total number of cells = 165),

F551C: n=4 (fields of view, total number of cells = 108),

C552: n=3 (fields of view, total number of cells = 90).

**Fig. S3G: DTT-stressed cells, area and intensity of Ire1-clusters:**

WT: n=395 cluster (raw data in Fig. 1D), cysteine-less: n=211 cluster (raw data in Fig. 1D), E540C: n=251 cluster, T541C: n=158 cluster, G542C: n=95 cluster, V543C: n=101 cluster, F544C: n=224 cluster, L545C: n=131 cluster, L546C: n=191 cluster, L547C: n=121 cluster, F548C: n=127 cluster, L549C: n=168 cluster, I550C: n=121 cluster, F551C: n=113 cluster, C552: n=75 cluster.

**Fig. S5A:** Δ*IRE1*, cysteine-less: n=12 (data from two individual colonies with technical replicates, identical with data in Fig. 5A), F531R F544C: n=12 (data from four individual colonies with technical replicates).

**Fig. S5B: DTT-stressed cells** F544C: n=14 (technical duplicates from seven individual colonies – identical with data in Fig. 3F), F531R F544C: n=4 (technical duplicates from two individual colonies).

**TM-stressed cells** F544C n=10 (technical duplicates from five individual colonies – identical with data in Fig. 3F), F531R F544C: n=4 (technical duplicates from two individual colonies).

**Fig. S6A:** WT: n=6 (technical replicates from two individual colonies); IF1: n=4 (technical replicates); IF2: n=12 (technical replicates from two individual colonies); Δ*IRE1*: n=6 (technical replicates).

**Fig. S6B:***reused for analysis*: E540C: n=15 (fields of view, identical with data in Fig. S3E) with the total number of cells = 374; T541C: n=9 (fields of view, identical with data in Fig. S3E) with the total number of cells = 181; F544C: n=19 (fields of view, identical with data in Fig. S3E) with the total number of cells = 440. *Additional data*: E540C: n=7 (fields of view, total number of cells = 281), T541C: n=6 (fields of view, total number of cells = 172), F544C: n=6 (fields of view, total number of cells = 213), E540C/IF2: n=7 (fields of view, total number of cells = 98), T541C/IF2: n=6 (fields of view, total number of cells = 150), F544C/IF2: n=6 (fields of view, total number of cells = 208).

---

**Fig. S1.**
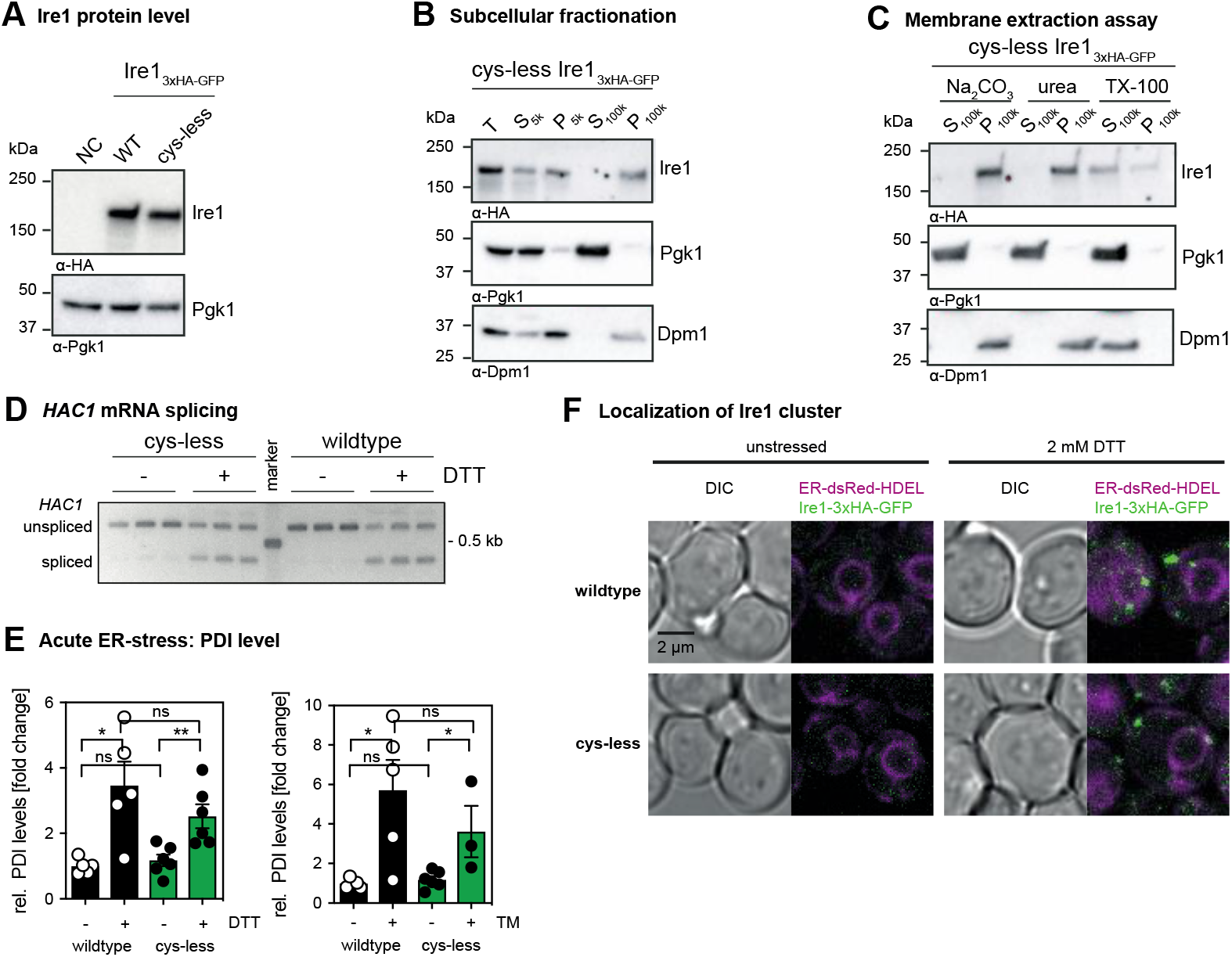
Protein levels of cysteine-less Ire1 and characterization of its membrane association. (*A*) Protein levels of cells expressing either *IRE1*_3xHA-GFP_ WT or the cysteine-less (cysteine-less) variant. The isogenic wildtype strain BY4741 that does not express a HA-tagged variant of IRE1 was used as a specificity control (NC). Stationary overnight cultures were used to inoculate a fresh culture in SCD complete to an OD_600_ of 0.2 and cultivated until an OD_600_ of 1 was reached. 0.1 OD equivalents of cell lysates were immunoblotted using anti-HA and anti-Pgk1 antibodies. (*B*) Subcellular fractionation of exponentially growing cells expressing cysteine-less *IRE1*_3xHA-GFP_ by differential centrifugation at 5,000 x g and 100,000 x g. Stationary overnight cultures were used to inoculate a fresh culture in SCD complete to an OD_600_ of 0.2 and cultivated until an OD_600_ of 1 was reached. 80 OD_600_ equivalents were harvested and used for microsomal membrane preparation. The individual supernatant and pellet fractions were analyzed by immunoblotting using anti-HA, anti-Pgk1 and anti-Dpm1 antibodies by loading 0.4 OD equivalents. (*C*) Extraction assay of microsomes. Carbonate and urea extraction validate proper membrane integration of cysteine-less *IRE1*_3xHA-GFP_ (cysteine-less). Samples of each step corresponding to 0.2 OD equivalents were analyzed by immunoblotting using anti-HA, anti-Pgk1 and anti-Dpm1 antibodies. (*D*) The indicated strains from a stationary culture were used to inoculate fresh culture in SCD to an OD_600_ of 0.2. After cultivation at 30°C to an OD_600_ of 0.7, cells were either left untreated or stressed with DTT (1 h, 2 mM, SCD). The level of the cDNA obtained from the spliced and unspliced *HAC1* mRNA was amplified and separated by a 2% agarose gel. (*E*) *PDI1* mRNA levels in acutely stressed cells normalized to the fold change of unstressed cells expressing *IRE1*_3xHA-GFP_ wildtype. Exponentially growing cells of the indicated strains were used to inoculated fresh YPD media to an OD_600_ of 0.2, cultivated in YPD and acutely stressed with either 4 mM DTT (left panel) or 1.0 μg/ml Tunicamycin (right panel) for 1 h. The relative level of *PDI1* in these cells was analyzed by RT-qPCR and quantitated using the comparative ΔΔCT method using normalization to *ACT1* levels. The data were normalized to the *PDI1* level in unstressed cells carrying the *IRE1*_3xHA-GFP_ wildtype construct. All error bars in this figure represent the mean ± SEM of at least three independent experiments. Significance was tested by an unpaired, two-tailed Student’s t test. **p<0.01, *p<0.05. (*F*) Cells were cultivated from OD_600_ of 0.2 to OD_600_ of 0.7 in SCD medium and then either left untreated or stressed with 2 mM DTT for 1 h. Life cells were mounted on agar slides and z-stacks were recorded using confocal microscopy. Images show the center plane of indicated channels.

**Fig. S2.**
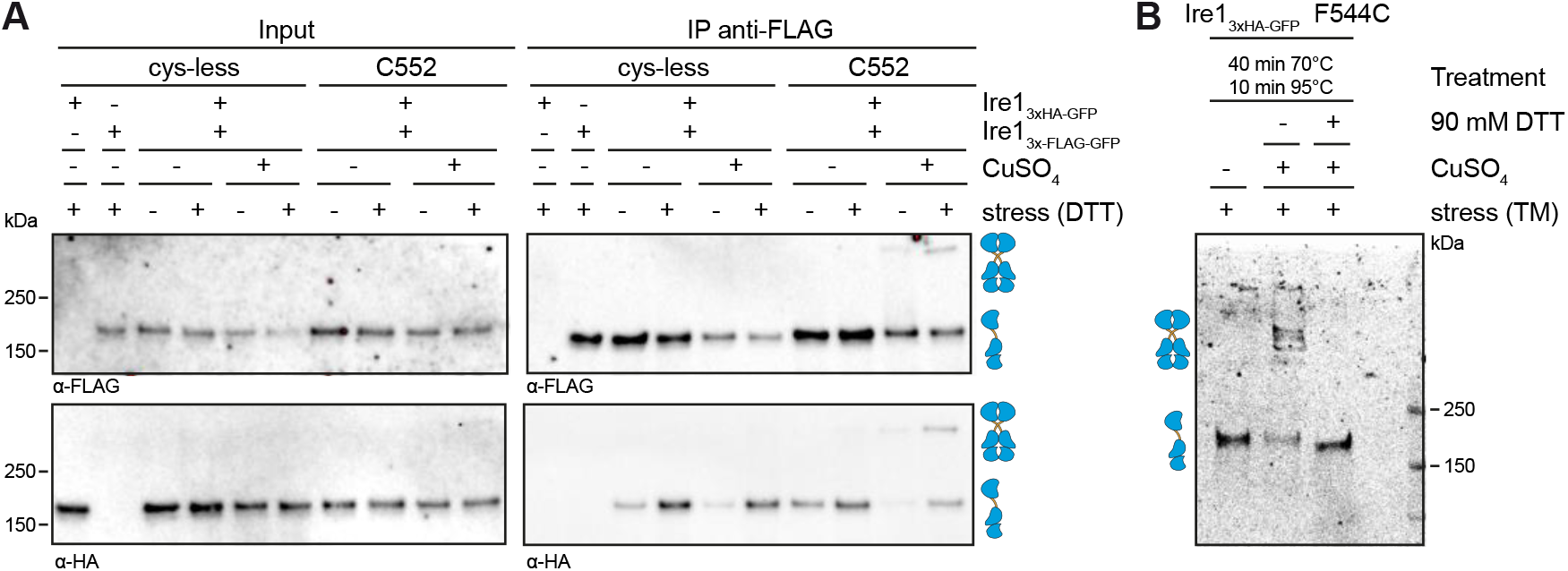
Validation of a covalent, reversible crosslinking of Ire1 homo-dimers via disulfide bridges. (A) A crosslinking experiment using CuSO_4_ was performed with microsomes prepared from cells expressing a HA-tagged variant of Ire1 from endogenous locus (*IRE1*_3xHA-GFP_) and a Flag-tagged variant (*IRE1*_3xFlag-GFP_) from a *CEN*-based plasmid. A yeast culture in selective SCD-LEU was inoculated to an OD_600_ of 0.2 from a stationary overnight culture and cultivated at 30°C until an OD_600_ of 0.7 was reached. The cells were either stressed with 2 mM DTT or left untreated and were further cultivated for 1 h. 80 OD_600_ equivalents from these cultures were harvested by centrifugation. Microsomal membranes were isolated by differential centrifugation. Microsomes prepared from cells expressing only one of the two tagged variants of Ire1 served as controls. Both constructs contained a single cysteine in the TMH region at the position 552 (C552). After incubation of the microsomes with 10 mM CuSO_4_ on ice for 5 min, the crosslinking reaction was stopped by the addition of NEM in a final concentration of 111 mM and EDTA in a final concentration of 50 mM. The microsomes were then solubilized using 2% Triton X-100 and subjected to an IP using anti-Flag beads. Both the input and IP samples were analyzed by immunoblotting using anti-Flag and anti-HA antibodies. (B) The reversibility of the cysteine-mediated crosslink was validated using the indicated F544C variant of Ire1_3xHA-GFP_. The crosslink was induced by CuSO_4_ in microsomes prepared from cells stressed with TM as described in Fig. 3. The crosslink was reverted by treating the sample with 90 mM DTT and incubating at 70° and 95° as indicated. The monomeric and dimeric species of Ire1 is indicated by symbols.

**Fig. S3.**
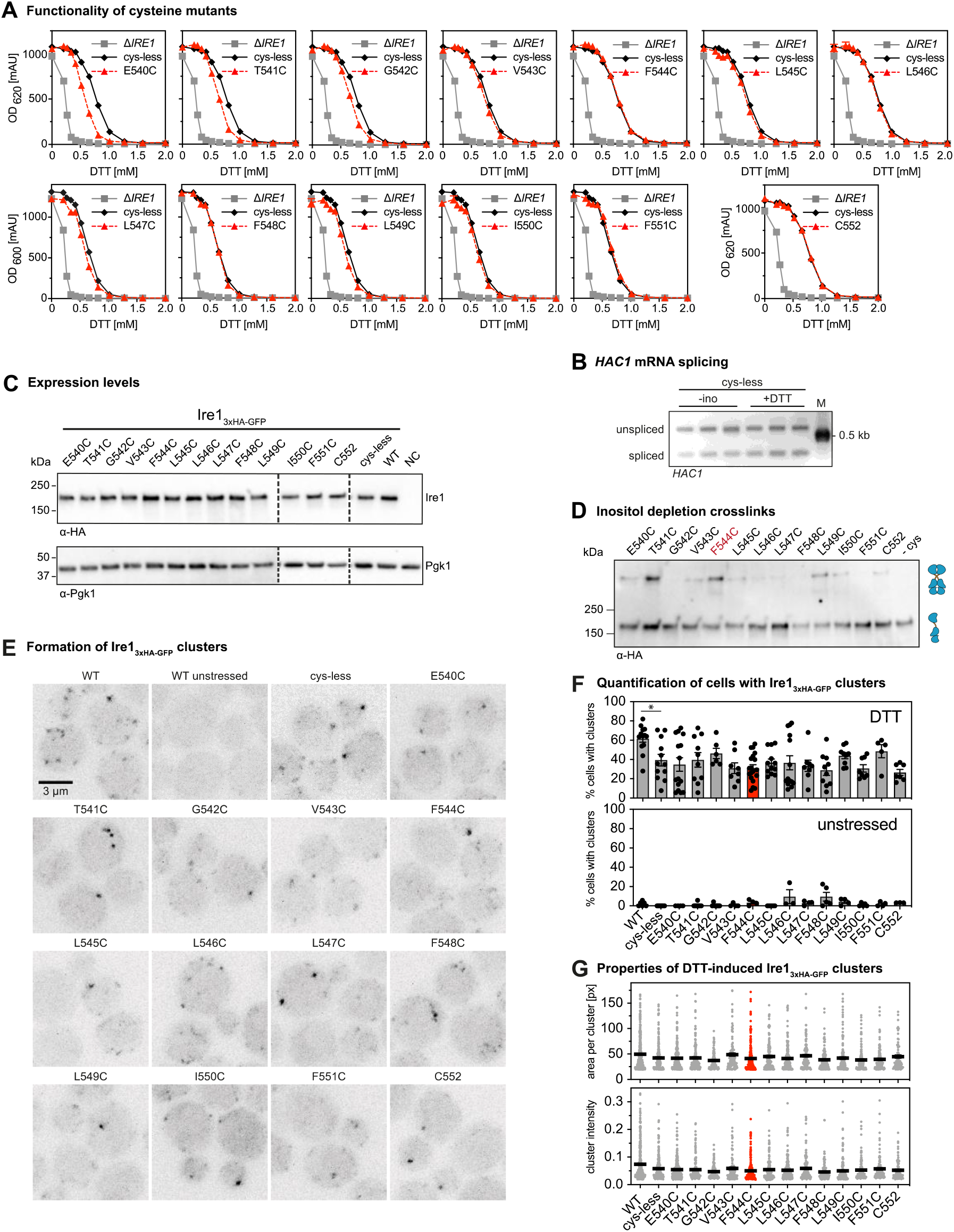
Functionality of cysteine mutants and their crosslinking potential in lipid bilayer stress conditions. (*A*) The resistance to ER-stress was investigated for the indicated yeast strains. Stationary overnight cultures of the indicated yeast strains were used to inoculate a fresh culture in full or minimal media to an OD_600_ of 0.2. After cultivation for 5 to 7 h at 30°C the cells were diluted with fresh minimal media to an OD_600_ of 0.1. Cells were cultivated for 18 h at 30°C and stressed with DTT. The density of the resulting culture was determined using the OD_620_ or OD_600_. The error bars represent the mean ± SEM of at least two independent clones. (*B*) The indicated strains were cultivated and treated as described in Fig. 3C and 3D using conditions of proteotoxic and lipid bilayer stress, respectively. The level of the cDNA obtained from the spliced and unspliced *HAC1* mRNA was amplified and separated by a 2% agarose gel. (*C*) Protein levels of cells expressing different *IRE1*_3xHA-GFP_ variants. The lysates of exponentially growing cells were immunoblotted using anti-HA and anti-Pgk1 antibodies. (*D*) Crosslinking of single cysteine variants of Ire1 in microsomes derived from cells grown in lipid bilayer stress conditions. Exponentially growing cells in SCD complete media were washed and used to inoculate a fresh culture in SCD complete to an OD_600_ of 0.5. To induce lipid bilayer stress, the cells were washed and then cultivated in pre-warmed SCD complete w/o inositol medium for 3 h. 80 OD equivalents were harvested and used for microsomal membrane preparation. CuSO_4_ induced crosslink was performed by incubating 8 μl of microsomes with 2 μl of 50 mM CuSO_4_ for 5 min on ice. After stopping the reaction with NEM and EDTA, samples were subjected to SDS-PAGE with a subsequent immunoblotting with anti-HA antibody. Notably, all samples subjected to SDS-PAGE underwent a crosslinking procedure. Differences in specific and unspecific crosslinking may falsely suggest differences in loading. (*E*) Cells were cultivated to the early exponential phase in SCD and either treated with 2 mM DTT for 1 h or left untreated. Representative images (maximum projections of z-stacks) recorded by confocal microscopy. (*F*) The percentage of cluster-containing cells was determined for stressed (2 mM DTT, 1 h) and unstressed cells using a custom-made CellProfiler pipeline (3). The percentage of cluster-containing cells with the cysteine-less variant of Ire1 is not significantly different from any of the cells with single-cysteine variants. The data for the wildtype variant of Ire1 and cysteine-less Ire1 are identical with the data from Fig. 1C and plotted as a reference. (*G*) The area of the detected clusters in the z-projection was determined and plotted. It was 49.9 px for the wildtype variant, 42.6 px for the cysteine-less variant, and ranged from a minimum of 37.2 px (G542C) to maximum of 48.9 px (V543C) for the single-cysteine variants. The integrated fluorescent intensity of detected clusters was 0.074 (arbitrary units) for the wildtype, 0.059 for the cysteine-less construct and ranged from a minimum of 0.046 for the F548C variant to a maximum of 0.059 for the L547C variant. *p<0.05.

**Fig. S4.**
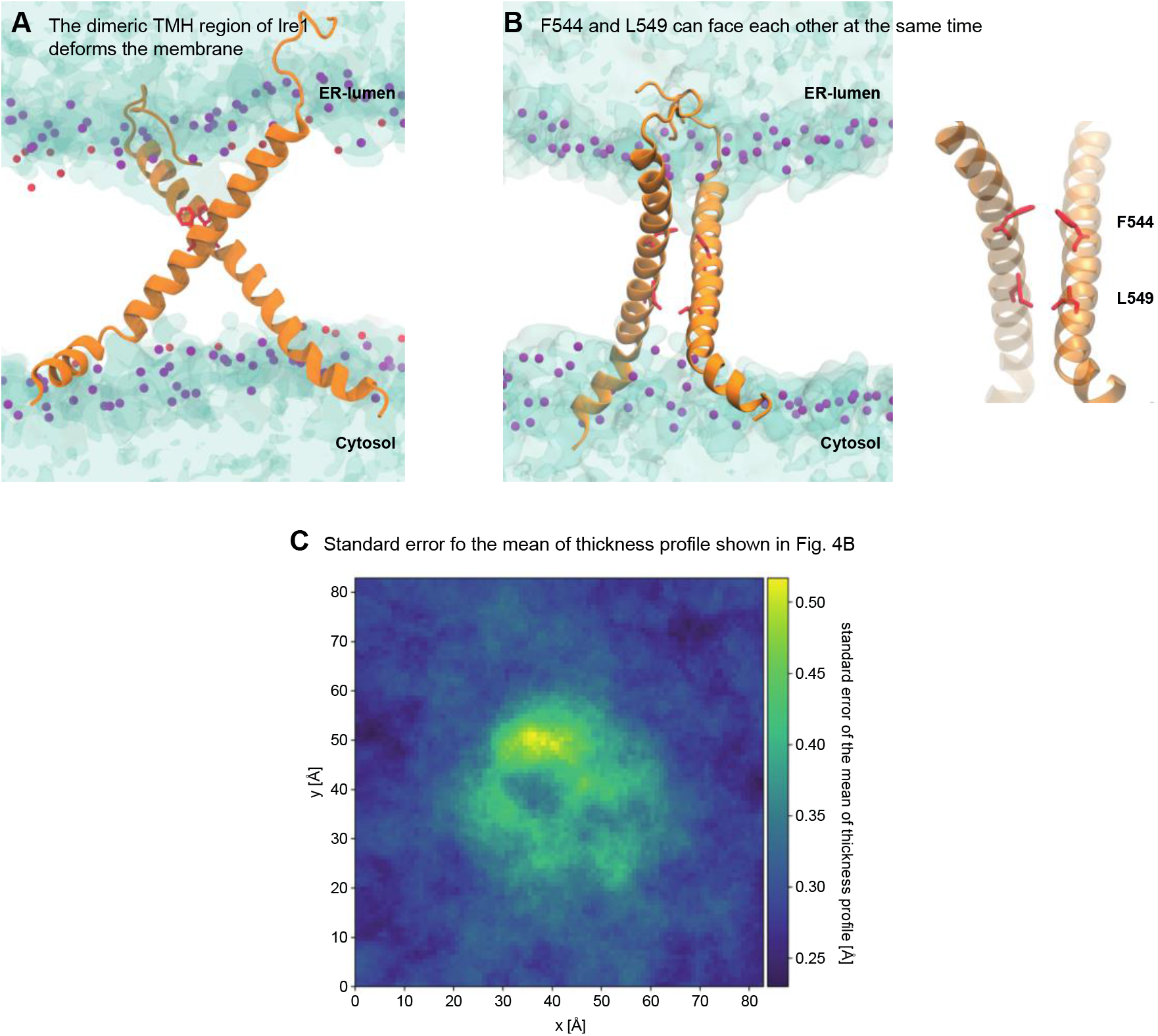
The dimeric TMH region of Ire1 deforms the membrane. (*A*) Membrane deformation by the modeled, dimeric TMH region of Ire1. Water is shown in blue tones with a transparent surface representation. The phosphate moieties of POPC are shown as purple beads. (*B*) Configuration of a model TMH dimer obtained from atomistic molecular dynamics simulations. Protomers are shown as an orange ribbon, with the residues F544 and L549 highlighted in red. The phosphate moieties of POPC are shown as purple beads. The hydroxyl groups of cholesterol molecules are shown as red beads. Water is shown with a transparent surface representation. In the right panel, lipid and water are not shown for clarity. (*C*) The standard error of the mean of the thickness profile represented in Fig. 4B. The thickness fluctuations in the close proximity of the TMH dimer (not shown, centered in the middle of the box) gives rise to a locally increased standard error of the mean of the thickness profile, but is much lower than the actual degree of membrane deformation as plotted in Fig. 4B.

**Fig. S5.**
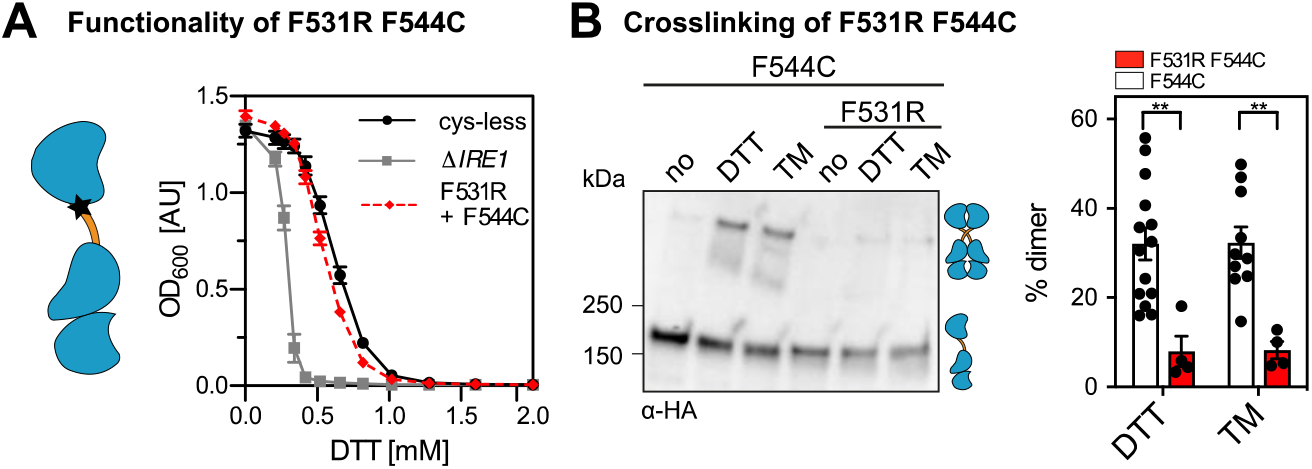
A mutation of the AH affects Ire1 function and crosslinking propensity. (*A*) The ER-stress resistance of cells expressing the AH-disrupting F531R variant of Ire1_3xHA-GFP_ containing the F544C single-cysteine was scored using an ER-stress resistance assay. The indicated cells were cultivated and treated as in Fig. 5A, C. (*B*) The impact of the AH-disrupting F531R mutation of Ire1 on the degree of crosslinking via the single-cysteine variant F544C was determined using the microsome-based crosslinking assay. Cells were cultivated and further treated as described in Fig. 5B, D. The data are represented as the mean ± SEM and are derived from at least three independent experiments. Significance was tested by an unpaired, two-tailed Student’s t test. **p<0.01.

**Fig. S6.**
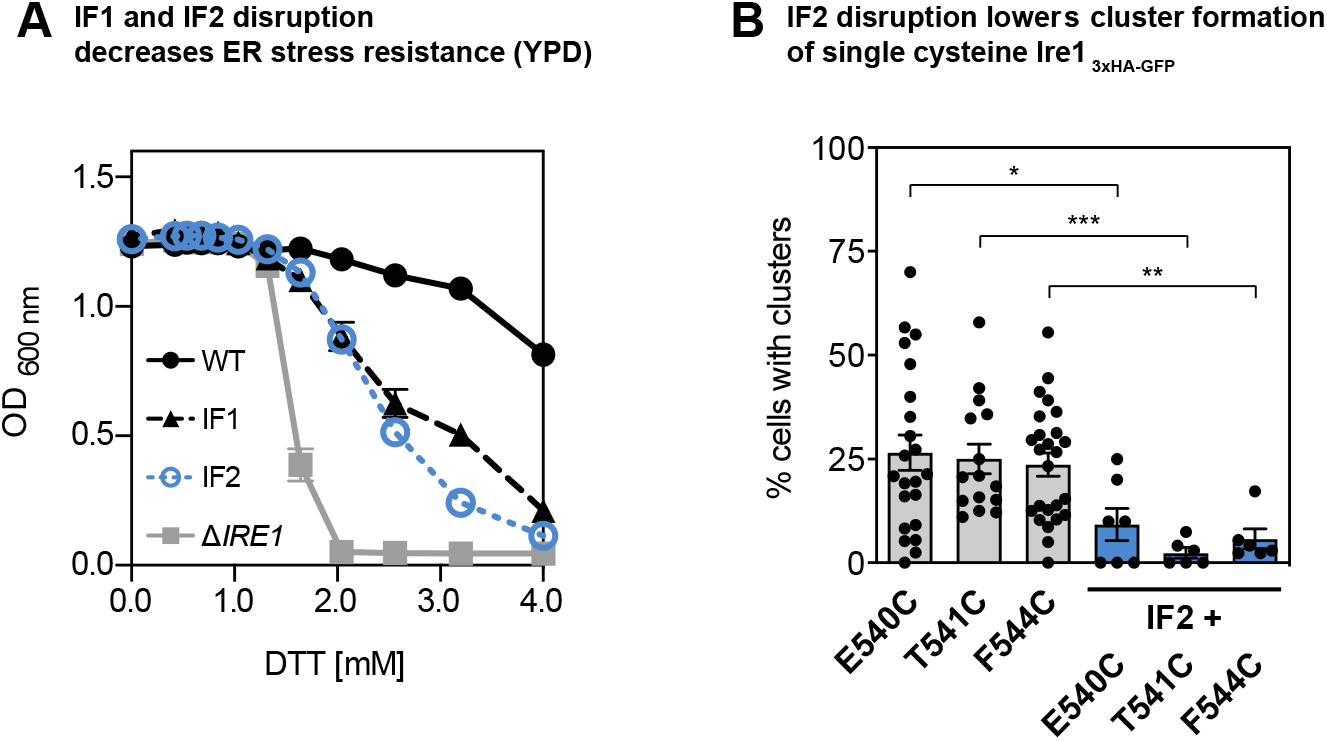
Disrupting ER-luminal interfaces for dimerization (IF1) and oligomerization (IF2) of IRe1 impairs cellular ER-stress resistance and the formation of Ire1 clusters. (*A*) The ER-stress resistance of cells expressing IF1 (T226A/F247A)- or IF2 (W426A)-disrupting knock-in variant of *IRE1* encoding all native cysteines was analyzed in rich medium. The indicated cells were cultivated and treated as in Fig. 5A, C. The data for reference strains are derived from cells lacking *IRE1* (gray) and from cells with a knock-in variant of *IRE1* containing all native cysteines (RE425). (*B*) The percentage of cluster-containing cells was determined for the indicated strains cultivated in SCD medium and stressed with 2 mM DTT for 1 h. Microscopic images were analyzed using a customized CellProfiler pipeline (3). The percentage of cluster-containing cells with single cysteine variants of Ire1 is significantly different from any of the cells where the ER-luminal IF2 was disrupted by mutation (W426A). The data are represented as the mean ± SEM. *p<0.05, **p<0.01, ***p<0.001.

**Table S1.** Yeast strains of used in this study.

| Strain | Description | Genotype | Source |
| --- | --- | --- | --- |
| RE001 | BY4741 | BY4741 MATa; *his3Δ1*; *leu2Δ0*; *met15Δ0*; *ura3Δ0* | Euroscarf |
| RE046 | $\Delta IRE1$ | BY4741 MATa; *his3Δ1*; *leu2Δ0*; *met15Δ0*; *ura3Δ0*; *ire1Δ::kanMX4* | Euroscarf |
| RE127 | $\Delta IRE1 \Delta IRE1$promotor | BY4741 MATa; *his3Δ1*; *leu2Δ0*; *met15Δ0*; *ura3Δ0*; *ire1Δ::URA* pUG72 | Halbleib *et al.* |
| RE425 | *IRE1*-3xHA-yeGFP | BY4741 MATa; *his3Δ1*; *leu2Δ0*; *met15Δ0*; *ura3Δ0*; *ire1Δ::URA* *IRE1*-3xHA-yeGFP::HIS pRE451 | Halbleib *et al.* |
| RE343 | *IRE1*-3xHA-yeGFP<br>*cysteine-less* | BY4741 MATa; *his3Δ1*; *leu2Δ0*; *met15Δ0*; *ura3Δ0*; *ire1Δ::URA* *IRE1*-3xHA-yeGFP::HIS pRE375 | This paper |
| RE342 | *IRE1*-3xHA-yeGFP C552 | BY4741 MATa; *his3Δ1*; *leu2Δ0*; *met15Δ0*; *ura3Δ0*; *ire1Δ::URA* *IRE1*-3xHA-yeGFP::HIS pRE374 | This paper |
| RE428 | *IRE1*-3xHA-yeGFP<br>W426A (IF2) | BY4741 MATa; *his3Δ1*; *leu2Δ0*; *met15Δ0*; *ura3Δ0*; *ire1Δ::URA* *IRE1*-3xHA-yeGFP::HIS pRE455 | Halbleib *et al.* |
| RE438 | *IRE1*-3xHA-yeGFP<br>T226A/F247A (IF1) | BY4741 MATa; *his3Δ1*; *leu2Δ0*; *met15Δ0*; *ura3Δ0*; *ire1Δ::URA* *IRE1*-3xHA-yeGFP::HIS pRE465 | Halbleib *et al.* |
| RE725 | *IRE1*-3xHA-yeGFP<br>*cysteine-less*<br>+ CEN *IRE1*-3xFLAG-yeGFP<br>*cysteine-less* | BY4741 MATa; *his3Δ1*; *leu2Δ0*; *met15Δ0*; *ura3Δ0*; *ire1Δ::URA* *IRE1*-3xHA-yeGFP::HIS pRE375 *IRE1*-3xFLAG-yeGFP::LEU pRE699 | This paper |
| RE726 | *IRE1*-3xHA-yeGFP C552 +<br>CEN *IRE1*-3xFLAG-yeGFP<br>C552 | BY4741 MATa; *his3Δ1*; *leu2Δ0*; *met15Δ0*; *ura3Δ0*; *ire1Δ::URA* *IRE1*-3xHA-yeGFP::HIS pRE374 *IRE1*-3xFLAG-yeGFP::LEU pRE700 | This paper |
| RE530 | *IRE1*-3xHA-yeGFP E540C<br>*single cysteine* | BY4741 MATa; *his3Δ1*; *leu2Δ0*; *met15Δ0*; *ura3Δ0*; *ire1Δ::URA* *IRE1*-3xHA-yeGFP::HIS pRE575 | This paper |
| RE531 | *IRE1*-3xHA-yeGFP T541C<br>*single cysteine* | BY4741 MATa; *his3Δ1*; *leu2Δ0*; *met15Δ0*; *ura3Δ0*; *ire1Δ::URA* *IRE1*-3xHA-yeGFP::HIS pRE576 | This paper |
| RE532 | *IRE1*-3xHA-yeGFP G542C<br>*single cysteine* | BY4741 MATa; *his3Δ1*; *leu2Δ0*; *met15Δ0*; *ura3Δ0*; *ire1Δ::URA* *IRE1*-3xHA-yeGFP::HIS pRE577 | This paper |
| RE533 | *IRE1*-3xHA-yeGFP V543C<br>*single cysteine* | BY4741 MATa; *his3Δ1*; *leu2Δ0*; *met15Δ0*; *ura3Δ0*; *ire1Δ::URA* *IRE1*-3xHA-yeGFP::HIS pRE578 | This paper |
| RE534 | *IRE1*-3xHA-yeGFP F544C<br>*single cysteine* | BY4741 MATa; *his3Δ1*; *leu2Δ0*; *met15Δ0*; *ura3Δ0*; *ire1Δ::URA* *IRE1*-3xHA-yeGFP::HIS pRE579 | This paper |
| RE522 | *IRE1*-3xHA-yeGFP L545C<br>*single cysteine* | BY4741 MATa; *his3Δ1*; *leu2Δ0*; *met15Δ0*; *ura3Δ0*; *ire1Δ::URA* *IRE1*-3xHA-yeGFP::HIS pRE570 | This paper |
| RE535 | *IRE1*-3xHA-yeGFP L546C<br>*single cysteine* | BY4741 MATa; *his3Δ1*; *leu2Δ0*; *met15Δ0*; *ura3Δ0*; *ire1Δ::URA* *IRE1*-3xHA-yeGFP::HIS pRE581 | This paper |
| RE717 | *IRE1*-3xHA-yeGFP L547C<br>*single cysteine* | BY4741 MATa; *his3Δ1*; *leu2Δ0*; *met15Δ0*; *ura3Δ0*; *ire1Δ::URA* *IRE1*-3xHA-yeGFP::HIS pRE691 | This paper |
| RE718 | *IRE1*-3xHA-yeGFP F548C<br>*single cysteine* | BY4741 MATa; *his3Δ1*; *leu2Δ0*; *met15Δ0*; *ura3Δ0*; *ire1Δ::URA* *IRE1*-3xHA-yeGFP::HIS pRE692 | This paper |
| RE719 | *IRE1*-3xHA-yeGFP L549C<br>*single cysteine* | BY4741 MATa; *his3Δ1*; *leu2Δ0*; *met15Δ0*; *ura3Δ0*; *ire1Δ::URA* *IRE1*-3xHA-yeGFP::HIS pRE693 | This paper |
| RE720 | *IRE1*-3xHA-yeGFP I550C<br>*single cysteine* | BY4741 MATa; *his3Δ1*; *leu2Δ0*; *met15Δ0*; *ura3Δ0*; *ire1Δ::URA* *IRE1*-3xHA-yeGFP::HIS pRE694 | This paper |
| RE721 | *IRE1*-3xHA-yeGFP F551C<br>*single cysteine* | BY4741 MATa; *his3Δ1*; *leu2Δ0*; *met15Δ0*; *ura3Δ0*; *ire1Δ::URA* *IRE1*-3xHA-yeGFP::HIS pRE695 | This paper |
| RE722 | *IRE1*-3xHA-yeGFP F544A<br>C552<br>*single cysteine* | BY4741 MATa; *his3Δ1*; *leu2Δ0*; *met15Δ0*; *ura3Δ0*; *ire1Δ::URA* *IRE1*-3xHA-yeGFP::HIS pRE696 | This paper |
| RE723 | *IRE1*-3xHA-yeGFP F531R<br>C552<br>*single cysteine* | BY4741 MATa; *his3Δ1*; *leu2Δ0*; *met15Δ0*; *ura3Δ0*; *ire1Δ::URA* *IRE1*-3xHA-yeGFP::HIS pRE698 | This paper |
| RE724 | *IRE1*-3xHA-yeGFP F531R<br>F544C<br>*single cysteine* | BY4741 MATa; *his3Δ1*; *leu2Δ0*; *met15Δ0*; *ura3Δ0*; *ire1Δ::URA* *IRE1*-3xHA-yeGFP::HIS pRE697 | This paper |
| RE773 | *IRE1*-3xHA-yeGFP W426A<br>E540C<br>*single cysteine* | BY4741 MATa; *his3Δ1*; *leu2Δ0*; *met15Δ0*; *ura3Δ0*; *ire1Δ::URA* *IRE1*-3xHA-yeGFP::HIS pRE575 | This paper |
| RE774 | *IRE1*-3xHA-yeGFP W426A<br>T541C<br>*single cysteine* | BY4741 MATa; *his3Δ1*; *leu2Δ0*; *met15Δ0*; *ura3Δ0*; *ire1Δ::URA* *IRE1*-3xHA-yeGFP::HIS pRE576 | This paper |
| RE776 | *IRE1*-3xHA-yeGFP W426A<br>F544C<br>*single cysteine* | BY4741 MATa; *his3Δ1*; *leu2Δ0*; *met15Δ0*; *ura3Δ0*; *ire1Δ::URA* *IRE1*-3xHA-yeGFP::HIS pRE579 | This paper |
| RE792 | *IRE1*-3xHA-yeGFP ER-dsRed-HDEL | BY4741 MATa; *his3Δ1*; *leu2Δ0*; *met15Δ0*; *ura3Δ0*; *ire1Δ::URA* *IRE1*-3xHA-yeGFP::HIS pRE451; *Kar2sig.seq.-dsRed-HDEL::natR* pRE850 | This paper |
| RE793 | *IRE1*-3xHA-yeGFP<br>*cysteine-less* ER-dsRed-HDEL | BY4741 MATa; *his3Δ1*; *leu2Δ0*; *met15Δ0*; *ura3Δ0*; *ire1Δ::URA* *IRE1*-3xHA-yeGFP::HIS pRE375; *Kar2sig.seq.-dsRed-HDEL::natR* pRE850 | This paper |

**Table S2.** Plasmids used in this study.

| Plasmid | Description | Recombinant DNA | Source |
| --- | --- | --- | --- |
| pRE451 | IRE1-3xHA-yeGFP | pcDNA3.1<br>IRE1-3xHA-yeGFP WT | Halbleib *et al.* |
| pEv200 | pRS315 IRE1-yeGFP-HA | pRS315 IRE1-yeGFP-HA | van Anken *et al.* |
| pRE375 | IRE1-3xHA-yeGFP<br>cysteine-less | pcDNA3.1 IRE1-3xHA-yeGFP<br>cysteine-less | This paper |
| pRE374 | IRE1-3xHA-yeGFP C552<br>single cysteine | pcDNA3.1 IRE1-3xHA-yeGFP C552<br>single cysteine | This paper |
| pRE455 | IRE1-3xHA-yeGFP W426A (IF2) | pcDNA3.1 IRE1-3xHA-yeGFP W426A (F2) | Halbleib *et al.* |
| pRE465 | IRE1-3xHA-yeGFP T226A/F247A (IF1) | pcDNA3.1 IRE1-3xHA-yeGFP<br>T226A/F247A (IF1) | Halbleib *et al.* |
| pRE699 | CEN IRE1-3xFLAG-yeGFP<br>cysteine-less | pRS315 IRE1-3xFLAG-yeGFP<br>cysteine-less | This paper |
| pRE700 | CEN IRE1-3xFLAG-yeGFP C552<br>single cysteine | pRS315 IRE1-3xFLAG-yeGFP C552<br>single cysteine | This paper |
| pRE575 | IRE1-3xHA-yeGFP E540C<br>single cysteine | pcDNA3.1 IRE1-3xHA-yeGFP E540C<br>single cysteine | This paper |
| pRE576 | IRE1-3xHA-yeGFP T541C<br>single cysteine | pcDNA3.1 IRE1-3xHA-yeGFP T541C<br>single cysteine | This paper |
| pRE577 | IRE1-3xHA-yeGFP G542C<br>single cysteine | pcDNA3.1 IRE1-3xHA-yeGFP G542C<br>single cysteine | This paper |
| pRE578 | IRE1-3xHA-yeGFP V543C<br>single cysteine | pcDNA3.1 IRE1-3xHA-yeGFP V543C<br>single cysteine | This paper |
| pRE579 | IRE1-3xHA-yeGFP F544C<br>single cysteine | pcDNA3.1 IRE1-3xHA-yeGFP F544C<br>single cysteine | This paper |
| pRE570 | IRE1-3xHA-yeGFP L545C<br>single cysteine | pcDNA3.1 IRE1-3xHA-yeGFP L545C<br>single cysteine | This paper |
| pRE581 | IRE1-3xHA-yeGFP L546C<br>single cysteine | pcDNA3.1 IRE1-3xHA-yeGFP L546C<br>single cysteine | This paper |
| pRE691 | IRE1-3xHA-yeGFP L547C<br>single cysteine | pcDNA3.1 IRE1-3xHA-yeGFP L547C<br>single cysteine | This paper |
| pRE692 | IRE1-3xHA-yeGFP F548C<br>single cysteine | pcDNA3.1 IRE1-3xHA-yeGFP F548C<br>single cysteine | This paper |
| pRE693 | IRE1-3xHA-yeGFP L549C<br>single cysteine | pcDNA3.1 IRE1-3xHA-yeGFP L549C<br>single cysteine | This paper |
| pRE694 | IRE1-3xHA-yeGFP I550C<br>single cysteine | pcDNA3.1 IRE1-3xHA-yeGFP I550C<br>single cysteine | This paper |
| pRE695 | IRE1-3xHA-yeGFP F551C<br>single cysteine | pcDNA3.1 IRE1-3xHA-yeGFP F551C<br>single cysteine | This paper |
| pRE696 | IRE1-3xHA-yeGFP F544A C552<br>single cysteine | pcDNA3.1 IRE1-3xHA-yeGFP F544A<br>C552 single cysteine | This paper |
| pRE697 | IRE1-3xHA-yeGFP F531R F544C<br>single cysteine | pcDNA3.1 IRE1-3xHA-yeGFP F531R<br>F544C single cysteine | This paper |
| pRE698 | IRE1-3xHA-yeGFP F531R C552<br>single cysteine | pcDNA3.1 IRE1-3xHA-yeGFP F531R<br>C552 single cysteine | This paper |
| pRE789 | IRE1-3xHA-yeGFP W426A E540C<br>single cysteine | pcDNA3.1 IRE1-3xHA-yeGFP W426A<br>E540C single cysteine | This paper |
| pRE790 | IRE1-3xHA-yeGFP W426A T541C<br>single cysteine | pcDNA3.1 IRE1-3xHA-yeGFP W426A<br>T541C single cysteine | This paper |
| pRE793 | IRE1-3xHA-yeGFP W426A F544C<br>single cysteine | pcDNA3.1 IRE1-3xHA-yeGFP W426A<br>F544C single cysteine | This paper |
| pSS455 | pRS303N TEF-Kar2signalsequence-dsRed-HDEL::natR (pRE850) | pRS303N TEF-Kar2signalsequence-dsRed-HDEL::natR | Sebastian Schuck<br>laboratory |

**Movie S1 (separate file). A structural model of the TMH region of Ire1 highlights membrane thinning and water penetration into the bilayer.** The two protomers of Ire1 TMH region are shown as orange ribbons. The residue corresponding to F544 is highlighted in red. The phosphate moieties of the lipid headgroups are shown as red/purple spheres. Water is indicated as shaded region to highlight membrane thinning.

**Movie S2 (separate file). Dynamics of the TMH region of Ire1 dimers over a period of 600 ns.** The two TMH regions are shown as orange ribbons. The residue corresponding to F544 is highlighted in red, while residues T541 to F551 are shown in blue. The phosphate moieties of the lipid headgroups are shown as purple spheres. Water, ions, and lipid acyl chains are omitted for clarity.

